# Characterization and decontamination of background noise in droplet-based single-cell protein expression data with DecontPro

**DOI:** 10.1101/2023.01.27.525964

**Authors:** Yuan Yin, Masanao Yajima, Joshua D. Campbell

## Abstract

Assays such as CITE-seq can measure the abundance of cell surface proteins on individual cells using antibody derived tags (ADTs). However, many ADTs have high levels of background noise that can obfuscate down-stream analyses. Using an exploratory analysis of PBMC datasets, we find that some droplets that were originally called “empty” due to low levels of RNA contained high levels of ADTs and likely corresponded to neutrophils. We identified a novel type of artifact in the empty droplets called a “spongelet” which has medium levels of ADT expression and is distinct from ambient noise. ADT expression levels in the spongelets correlate to ADT expression levels in the background peak of true cells in several datasets suggesting that they can contribute to background noise along with ambient ADTs. We then developed DecontPro, a novel Bayesian hierarchical model that can decontaminate ADT data by estimating and removing contamination from these sources. DecontPro outperforms other decontamination tools in removing aberrantly expressed ADTs while retaining native ADTs and in improving clustering specificity. Overall, these results suggest that identification of empty drops should be performed separately for RNA and ADT data and that DecontPro can be incorporated into CITE-seq workflows to improve the quality of downstream analyses.

## Introduction

Cellular indexing of transcriptomes and epitopes by sequencing (CITE-seq) is an assay that can quantify the abundance of RNA transcripts as well as cell surface proteins on individual cells^1^. Antibody-derived tags (ADTs) that bind to cell surface proteins are measured during sequencing to produce an ADT count matrix that quantifies the levels of these surface proteins. Information on the abundance of proteins complements single-cell RNA-seq (scRNA-seq) data and improves the ability to describe cell type and cell states with functional annotations^2–4^. Other variants of CITE-seq have been developed which can measure proteins in other settings such as CRISPR perturbations, single-cell ATAC-seq, or spatially resolved expression^5–7^.

Previous studies have noted that individual ADTs often have a multimodal distribution including one lower “background” peak and one or more higher peaks attributed to the true signal from the cells. The lower background peak has been attributed to noise from non-specific binding of antibodies^8^. Algorithms such as TotalVI^9^ have tried to leverage the multi-modal nature to identify and remove the lower background peak for each ADT. Other approaches try to measure and remove background levels using “spike-in” reference cells from another species such as mouse^1^. Utilizing spike-ins adds extra complexity to the experimental design and assumes that rate of contamination in the reference cells will be the same for the cells of interest in the dataset. Additionally, none of these approaches quantify specific sources of contamination within each cell.

Contamination from various sources can contribute to poor-quality data in single-cell assays. In scRNA-seq data, ambient RNA from the cell suspension can be counted along with a cell’s native RNA and result in contamination of gene markers between cell types^10,11^. Ambient contamination may also occur in CITE-seq data as the methods for generating CITE-seq data also rely on microfluidic droplet-based devices. Two computational methods, dsb^12^ and scAR^13^, have been proposed that use the ADT expression profiles of the “empty droplets”, i.e., droplets without a true cell to estimate and remove the noise from ambient material. However, these methods treat the empty droplets as a single source of noise. Furthermore, reliance on the empty droplets data may limit their application in cases where the empty droplet matrix is not available.

In this study, we analyzed four CITE-seq datasets and showed that there are at least four different types of droplets including 1) droplets containing true cells with high RNA and high ADT content; 2) droplets with low RNA content and high ADT content that are mislabeled as “empty droplets”; 3) droplets containing low levels ADTs matching ambient distributions; and 4) droplets containing medium levels of ADTs with non-specific distributions, which we denote as “spongelets”. We show that the ADT expression profiles of spongelets are highly correlated with the expression profiles of the background peak in true cells and likely contribute to contamination along with ambient ADTs. Based on these results, we developed a novel Bayesian hierarchical model called DecontPro (**Decont**amination of **Pro**tein expression data) that removes the background peak by estimating ambient contamination as well as contamination derived from other sources such as spongelets or non-specific binding. When applied to different ADT datasets, DecontPro was able to preserve the expression of native markers in known cell types while removing contamination from the non-native markers. DecontPro outperformed other tools in removing non-native markers and in improving downstream clustering in several benchmarking datasets. Finally, we show that DecontPro can increase the specificity of PD-1 expression in activated T and B-cells.

## Results

### Different contamination profiles contribute to CITE-seq data

We performed an exploratory analysis of CITE-seq and Total-seq datasets to understand heterogeneity among the droplets and to characterize different sources of contamination. We first analyzed a public dataset containing peripheral blood mononuclear cells from a healthy donor from 10x Genomics (PBMC 10K). Four distinct clusters of droplets were observed when comparing the total UMI counts of ADTs to total UMI counts of RNAs in each droplet (**Fig. 1A**). Cluster A had high counts for both RNA and ADTs and were called true cells by Cell Ranger (n=7,864 droplets). Clusters B-D were called empty droplets by Cell Ranger and had low RNA counts with varying levels of ADT counts. Cluster D contained droplets with low levels of both total RNA and total ADT counts (n=145,322 droplets). The average profile of the droplets from this cluster was highly correlated with the average profile of droplets from the Cell cluster for both RNA and ADTs for most datasets (R > 0.950) except for Golomb et al^14^ (R = 0.769; **Supplementary Fig. S1**) demonstrating that these droplets likely contain only ambient material^9^.

**Figure 1.**
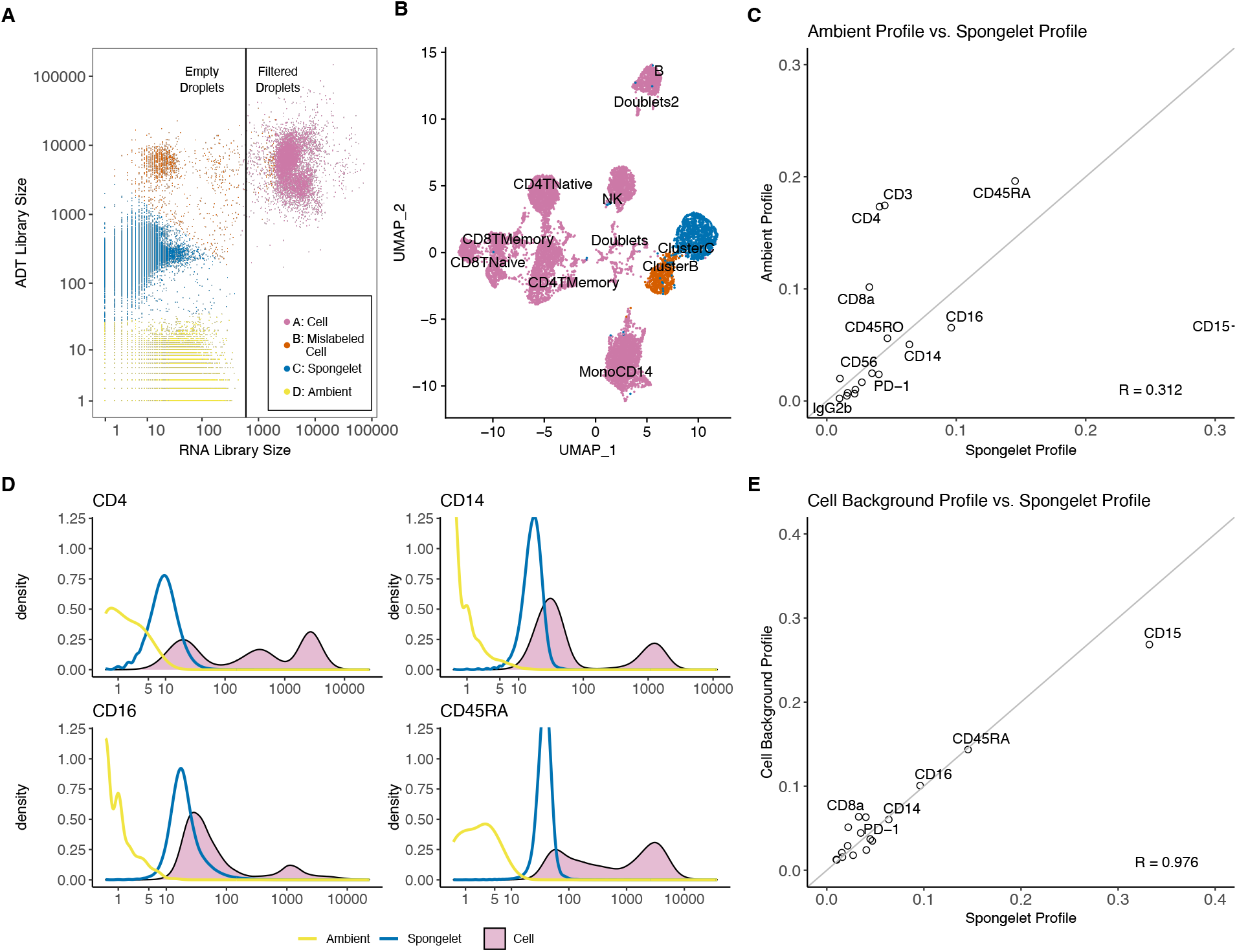
Analysis of ADT droplets in the PBMC 10K dataset. **A.** Among all droplets, only the cluster with high RNA-seq library size were filtered as cell droplets by the 10x Genomics software, while we observed clusters with varying levels of ADT library sizes among the droplets labeled as empty. We identified three empty droplet clusters, and called them clusters B, C and D. **B.** A thousand droplets sampled from clusters C and D each were combined with Cell droplets for clustering using Seurat workflow. Distinct clusters were observed. **C.** The empirical distribution built using the droplets from cluster C (spongelet) and D (ambient) showed poor correlation. **D.** Density plots of droplets from the cluster C were close to the background peak of the Cell density plot. **E.** We manually curate the background peaks of the Cell density and normalize them into a multinomial distribution. The distribution was found to be highly correlated with the spongelet cluster empirical distribution.

Cluster B had an average of 5,168 ADTs counts per droplet (n=1,406 droplets) while cluster C had an average of 268 ADTs (n=70,157 droplets). To understand the ADT profiles in these clusters, we randomly sampled a thousand droplets from each of these two clusters and analyzed them with the droplets containing cells from cluster A using the standard Seurat clustering workflow^15^. Droplets from the clusters B and C formed their own distinct clusters (**Fig. 1B**). Cluster B had significantly higher levels of CD15 and CD16 compared to other populations, but few differentially expressed genes in the RNA data (**Supplementary Fig. S2**). This “mislabeled cell” cluster likely represents neutrophils which are prevalent in white blood cells but have low RNA content^16,17^ and suggests that viable cells with low RNA content may be readily characterized by ADT expression. Therefore, filtering of empty droplets should be performed separately for ADT and RNA data. In contrast to the mislabeled cell cluster B, cluster C did not show strong enrichment for any ADTs. The average profile of droplets in cluster C was not highly correlated to the average profile of the ambient cluster D (R = 0.312; **Fig. 1C**), confirming that the source of this cluster is not solely related to ambient ADTs. We assigned the name “spongelets” to the droplets in cluster C given that they contain medium levels of ADTs that do not have enrichment in specific cell types.

The distribution of individual ADT expression in true cells is often multi-modal and contains more than one peak. For example, cells in cluster A have distributions of CD14, CD16, and CD45RA as bi-modal and the distribution of CD4 as tri-modal (**Fig. 1D**). The lower peak of the multi-modal densities has previously been characterized as background signals from non-specific binding of antibodies^8^. Interestingly, the density of ADTs in the spongelet cluster largely overlapped with the density of the lower peak in the cluster A (**Fig. 1D, Supplementary Fig. S3**). This overlap was also observed in the three other ADT datasets (**Supplementary Figs. S4 – S7**). The average percentage of each ADT in the lower peak of cluster A was highly correlated with the average percentage of each ADT in cluster C in the PBMC 10K dataset (R = 0.976; **Fig. 1E**), and to a lesser degree in two other datasets (R = 0.720 and R = 0.641; **Supplementary Fig. S8**). The low-level background expression indicated by the lower peak is also manifested after normalizing Cell droplets by their library sizes and classifying them by their cell types. (**Supplementary Fig. S9-S11**). Low peaks of ADTs were observed in cell types that are not supposed to express the ADTs. Interestingly, the density showing the background peak for each ADT has a consistent mean across cell types. Overall, these findings suggest an association between the ADT expression profiles of spongelets and the lower background peaks observed in true cells.

### A novel model for estimating and removing decontamination

We next sought to build a deconvolution algorithm that can estimate and remove contamination from the ambient material as well as any other sources contributing to the background including spongelets and non-specific binding. We assume that each cell is a mixture of three sources: 1) ADTs from the native cell, 2) ADTs from the ambient material present in the cell suspension, and 3) ADTs from any contamination source that contributes to a lower non-specific background peak (**Fig. 2A**).

**Figure 2.**
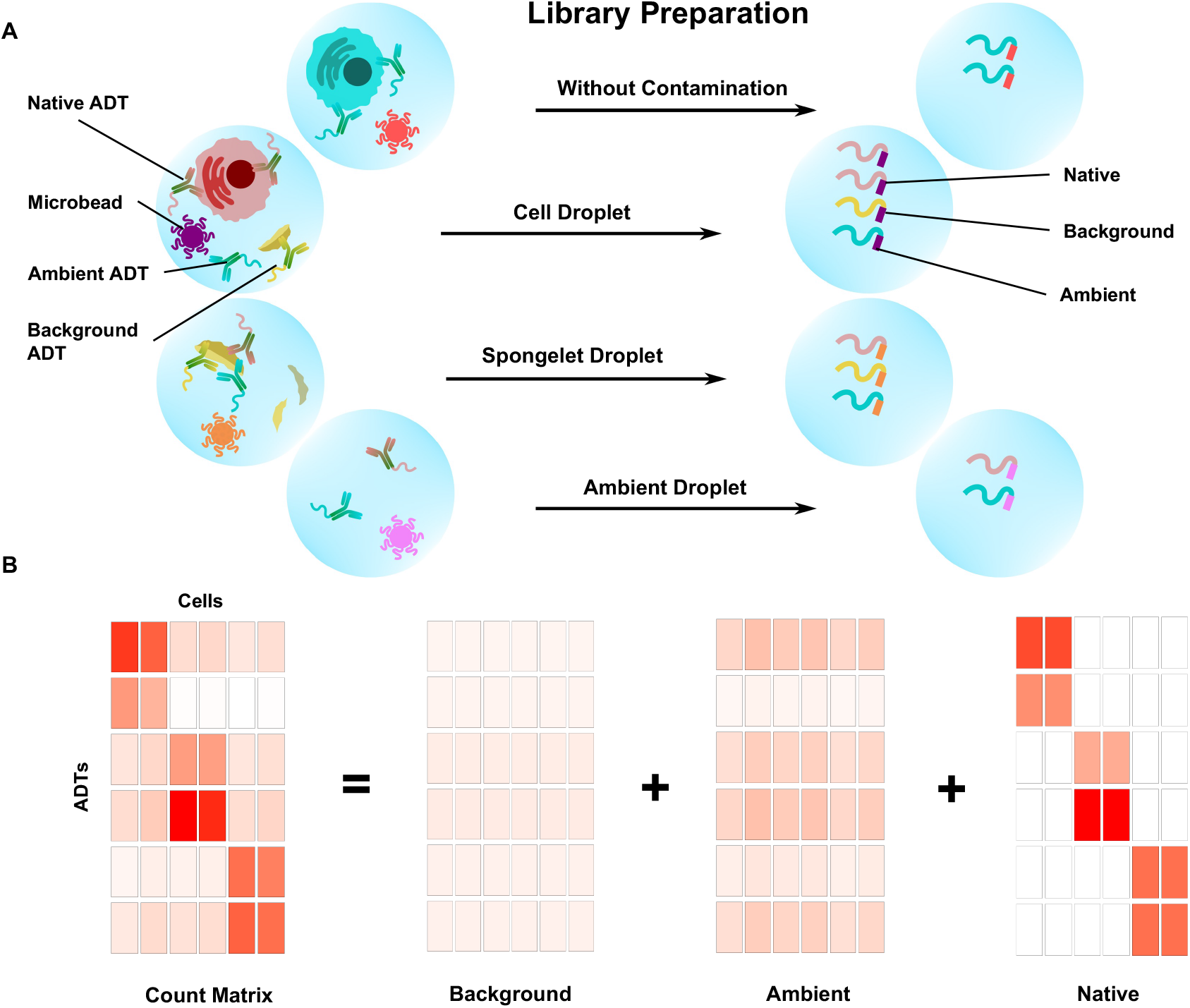
DecontPro can estimate and remove contamination from multiple sources in CITE-seq data. **A.** Single cells and ADTs are isolated in droplet compartments. Spongelet droplets contain some non-specific distribution and an intermediary level of ADTs, while ambient droplets have a low level of ADTs attributed to the ambient material from the cell suspension. Native ADTs refer to those that bind to their target cell surface proteins in the appropriate cell population. Background ADTs represent those from contamination sources other than ambient such as spongelets or non-specific binding of ADTs. These different sources of ADTs can be present and quantified together in each droplet during library preparation. **B.** The DecontPro model can deconvolute an ADT count matrix into a background matrix, an ambient matrix, and a native matrix along with the proportion of counts attributed to each source in each cell. The native matrix can be used in downstream analyses.

The native ADT expression of a cell population *k* is characterized by a multinomial distribution *φ_k_*, where *φ_kj_* represents the percentage of native expression attributed to ADT *j*. The ambient profile is characterized by a multinomial distribution *η* where *η_j_* represents the percentage of ambient expression attributed to ADT *j*. We use a mixing parameter *θ_ij_* to model the proportion of counts contributed by the ambient ADT *j* to the droplet *i*, and *β_ij_* to model the proportion of counts from other background sources for ADT *j* in droplet *i*. Parameter *μ_j_* is the prior for *β_ij_* and represents the average background level for ADT *j* in the dataset while parameter *δ_i_* is the prior for *θ_ij_* and represents the average ambient contamination level for droplet *i*. After mixing the parameters and scaling by the library size, the observed counts are generated from a Poisson distribution.

When the raw matrix with empty droplets is available, the ambient profile *η* can be estimated using the empirical distribution of the ambient droplets (i.e., droplets with low expression of ADTs). If the raw matrix is not provided, the ambient profile *η* for a dataset can be estimated by calculating the average of the ADT across filtered cell droplets. This is due to the fact that the average cell profile is highly correlated to the empirical distribution of the ambient droplets in the majority of datasets. For example, we observed this pattern in three ADT datasets (PBMC 10k: R = 0.977, PBMC 5k: R = 0.991, MALT 10k: R = 0.968; **Supplementary Fig. S1**). The correlation is less strong for one dataset (Golomb: R = 0.769), primarily due to an outlier ADT Ly6C. Ly6C had uniformly high level of expression across droplets while other highly expressed ADTs were more localized to specific clusters (**Supplementary Fig. S12**). We use the variational inference framework provided by Stan to estimate the remaining parameters and deconvolute the ADT count matrix into the Native, Ambient, and Background matrices for downstream analysis (**Fig. 2B**). Detailed description of the model can be found in the **Methods** section.

### Decontamination of PBMCs

To demonstrate the ability of our method to improve data quality, we applied DecontPro to the PBMC 10k dataset from 10x Genomics and compared the distributions of counts before and after decontamination. In the original count matrix, all ADTs were expressed to some degree in nearly every cell cluster (**Fig. 3A**). After decontamination, known markers for cell types remained highly expressed (**Fig. 3B**). For example, CD19 is a B-cell marker with an average expression of 668.46 in the B-cell cluster and a low average expression of 13.68 across other cell types in the original count matrix. After decontamination the only clusters retaining expression of CD19 were the B-cells and doublets. Similarly, T-cell marker CD3 had remaining counts only in the T-cell clusters and a doublet cluster.

**Figure 3.**
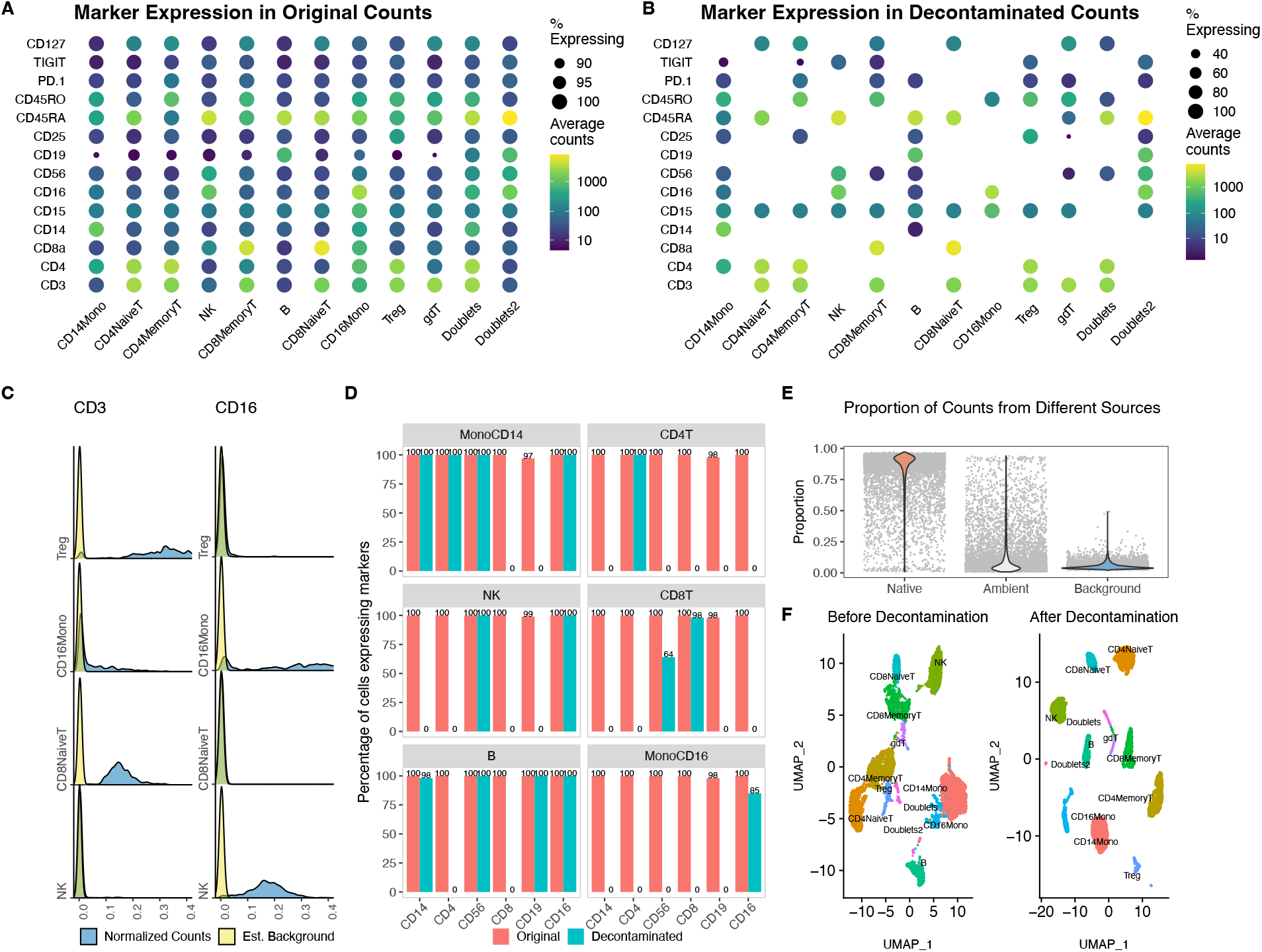
Decontamination of the PBMC 10k dataset with DecontPro. **A.** Expression profiles of ADTs in cell populations before decontamination. All ADTs were expressed in each cell population to some degree **B.** After decontamination with DecontPro, aberrantly expressed ADTs were removed or greatly reduced in non-native cell populations. **C.** The density of normalized ADT expression for CD3 and CD16 in selected cell populations. The model estimated background was superimposed onto the density plot. **D.** Percentage of cells expressing known cell type markers before and after decontamination in different cell populations. Markers with a count greater than one were considered expressed in a cell. **E.** The DecontPro model estimates the proportion of counts coming from native, ambient and background sources in each droplet. The median percentage of native, ambient, and background counts was 89.5%, 4.4%, and 5.5%, respectively. **F.** Using decontaminated counts improved separation of clusters on a UMAP.

For most markers, the density of the estimated background largely overlapped with the lower peak across cell clusters. For example, the lower levels of CD3 were estimated to be largely background and removed in the CD16 Monocyte and NK clusters (**Fig. 3C**). Similarly, for CD16 the lower levels of expression were estimated to be largely background in Naïve CD8 T-cells and T-regs clusters. As expected, native ADTs in their corresponding clusters were expressed at a higher level, and hence had a higher peak in the normalized counts’ density, such as CD3 in CD8 Naïve T-cells cluster and T-regs cluster.

To systematically assess how well the algorithm performed in specificity in retaining native ADTs for clusters, we calculated the percentage of cells in each cluster expressing native marker ADTs before and after decontamination (**Fig 3D**). Markers and cell types included CD3 and CD4 for CD4+ T-cells, CD3 and CD8 for CD8+ T-cells, CD19 for B-cells, CD14 for CD14+ monocytes, CD16 for CD16+ monocytes, and CD56 for NK cells. In many cases, the algorithm was able to greatly reduce or completely remove aberrant counts in non-native cell types. For example, CD14 expression was removed from T-cells, CD19 was removed from T-cells, NK-cells, and monocytes, and CD3 was removed from B-cells and monocytes. In some cases, the reduction left some ADT values in unexpected cell clusters. The monocyte marker CD14 was still detected in a high percentage of B-cells after decontamination (98%). However, the overall level of CD14 in B-cells was still greatly reduced compared to the original ADT counts (**Supplementary Fig. S13**).

DecontPro also estimates the percentage of counts contributed by the native, ambient and background signals (**Fig. 3E**). As expected, the native counts took the highest percentage of libraries across droplets on average (median 89.5%, range 1.0% - 97.1%), whereas the background counts were estimated at a consistently lower amount (median 4.4%, range 2.0% - 49.2%). The percentage of ambient ADTs had a similar median to the percentage of background counts, but a much larger range (median 5.5%, range 0.8% - 93.9%). Lastly, the clusters in the UMAP generated with the decontaminated counts were more separated compared to clusters in the UMAP generated with the original counts (**Fig. 3F**).

### Benchmarking against Other Methods

We benchmarked DecontPro against three other decontamination algorithms applicable to CITE-seq datasets: dsb, scAR, and totalVI. To compare how well decontamination improved clustering, we calculated the mean silhouette width of each cell cluster identified by Seurat before and after decontaminating using four public datasets. A higher mean silhouette width indicates higher similarities between cells within each cluster. Across all datasets, DecontPro showed the highest average improvement in silhouette widths after decontamination (**Fig. 4A**). We next sought to understand how effectively various algorithms improved the specificity of marker expressions in cell clusters. Specifically, we wanted to measure if the expected markers in annotated clusters retained their expression, while the unexpected markers from other cell types were removed. We calculated two scores for each algorithm on each dataset. A positive score was calculated by finding the percentage of cells in a cluster that express the expected “native” markers while a negative score calculated the percentage of cells in a cluster that express unexpected “nonnative” markers from other cell types. An effective decontamination algorithm will have a positive score close to 100 indicating that the expression of native markers was retained in their true cell population as well as a negative score close to 0 indicating that the expression of non-native markers from other cell types was successfully removed. The full list of annotated native markers for each cell type can be found in the **Methods** section. As a reference, the uncorrected original count data had high positive scores in all datasets and high negative scores close to 100 for the PBMC 10k, PBMC 5k and MALT 10k datasets, indicating a high level of contamination in most cell types (**Fig. 4B**). When using the difference between positive score and negative score to measure the algorithm performance, DecontPro showed high overall score, indicating that it retained the markers while removing the unexpected markers across datasets (PBMC 10k: 99, PBMC 5k: 99, MALT 10k: 98, Golomb: 97). As dsb performs its own normalization and scaling, the decontaminated count matrix is not directly comparable with the original count matrix. We therefore used two different thresholds for a maker to be considered detected in a cell (dsb: 1; dsb high threshold: 5) and calculated the scores accordingly. The higher threshold for dsb produced better negative scores more similar to algorithms such as DecontPro and scAR, at the expense of decreasing the positive scores in each dataset. The second-best performing algorithm across datasets based on difference between positive and negative scores was scAR (PBMC 10k: 96, PBMC 5k: 90, MALT 10k: 99, Golomb: 82).

**Figure 4.**
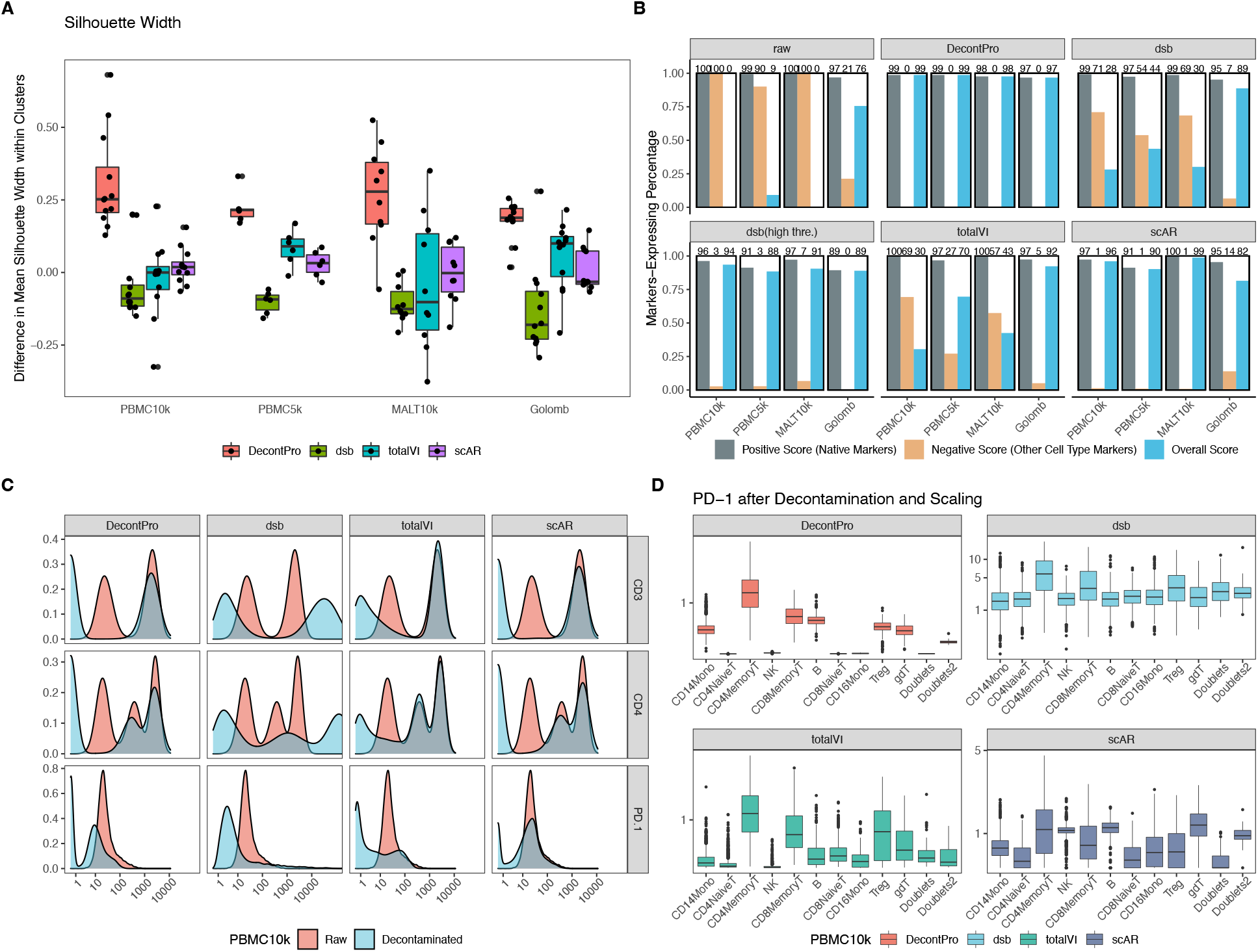
Benchmarking of ADT decontamination methods. We compared the ability of four algorithms to remove contamination from four datasets. The other methods include dsb, scAR, and totalVI. **A.** The average silhouette width for each cell cluster identified by Seurat was calculated before and after decontamination of ADT counts. A higher difference in silhouette widths indicates better improvement in cell similarity within that cluster. On average, DecontPro improved silhouette widths better than other methods for each dataset. **B.** For each dataset, a pair of scores were calculated after applying each decontamination algorithm to ascertain the degree to which each algorithm maintains true expression while removing contamination. The “positive” score was defined as the percentage of cells expressing native markers averaged across cell clusters. In contrast, the “negative” score was defined as the percentage of cells expressing non-native markers averaged across cell clusters. For example, in the CD4 naïve T-cells cluster, CD4 and CD45RA were defined as the native markers, while CD8 and CD45RO were defined as the non-native markers. As dsb outputs normalized continuous counts after decontamination, we applied two different thresholds to determine which markers were detected in each cell. The overall score is the positive score minus the negative score. Overall, DecontPro performs the best in the overall score in all four datasets. **C.** Density plot of CD3, CD4 and PD-1 before and after decontamination in the PBMC 10k dataset. The dsb output was exponentiated to allow for comparison with the uncorrected original count density. **D.** Box plot of PD-1 level in cell clusters after decontamination and scaling PBMC 10k dataset.

**Figure 5.**
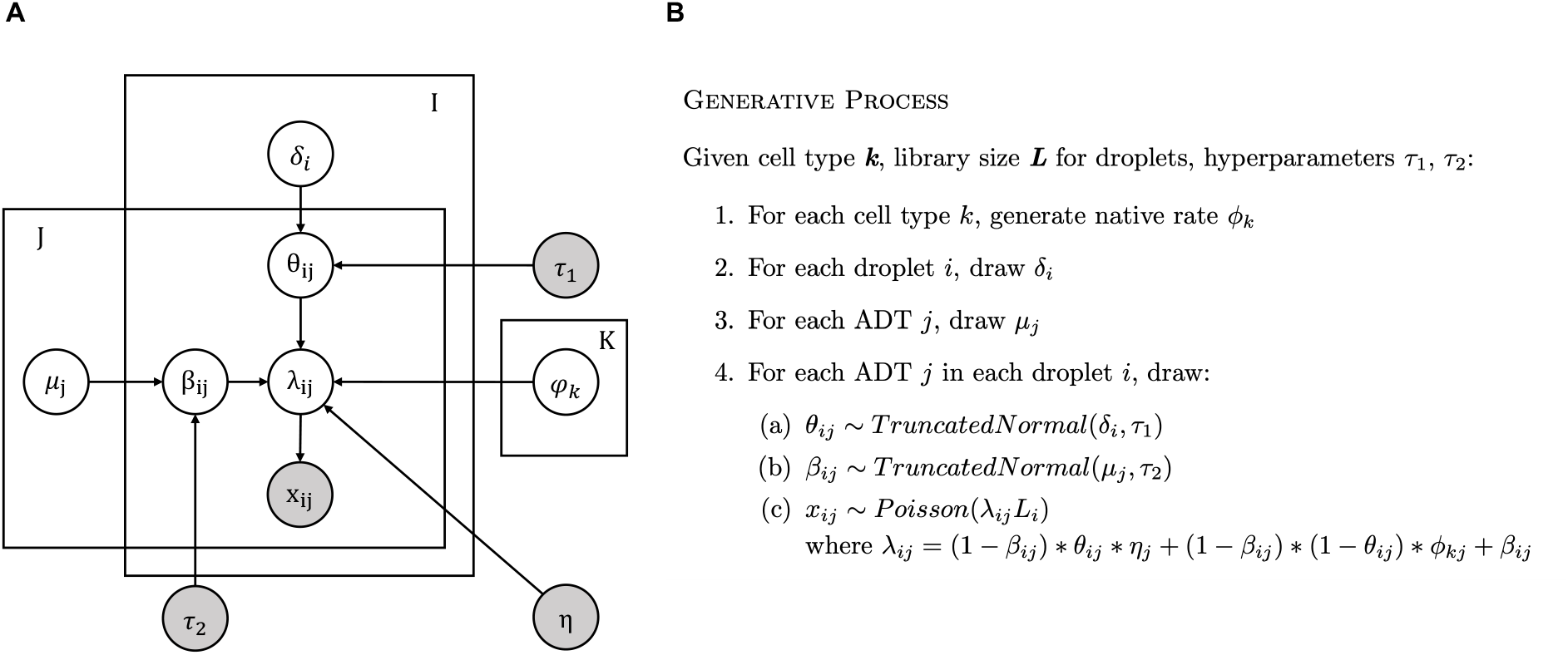
Summary of the DecontPro model. **A.** Plate diagram for the DecontPro model. **B.** The generative process of the DecontPro model.

For the PBMC 10k dataset, the decontamination results for known cell type markers were mostly similar across algorithms. For antibodies with high expression, such as CD3 and CD4, all algorithms retained the higher peak while significantly reducing the lower background peak (**Fig. 4C**). However, PD-1 was expressed at lower levels and demonstrated a noticeable difference in decontamination results between algorithms. PD-1 is known to be predominantly expressed in activated T-cells, along with B-cells, natural killer (NK) cells and myeloid cells^18–20^. Our algorithm reduced the PD-1 expression significantly in the naïve T-cells population, while preserving the expression level in the CD4 memory T-cells, CD8 memory T-cells, and Treg clusters where we expected a high PD-1 level (**Fig. 4D**). Additionally, some levels in the B-cells and monocytes clusters were also retained. Dsb and totalVI largely repressed PD-1 expression in all clusters except CD4 memory T-cells, CD8 memory T-cells, and Treg cluster. ScAR is the only algorithm that retained PD-1 in the NK cells cluster but has also removed its expression in the Treg cluster.

## Discussion

We developed DecontPro, a Bayesian statistical model, that decontaminates two sources of contaminations we observed empirically in CITE-seq data. Previously, the empty drops were thought to only contain noise from ambient material from the cell suspension. Our analyses suggest that the empty droplets filtered using RNA are heterogeneous and may contain different sources of signal and noise. One cluster of empty droplets had high levels of specific ADTs and thus likely contained true cells. These results suggest the need for empty drop calling to be performed separately for ADT and RNA data. We also identified a cluster of empty droplets with medium levels of ADTs containing a different profile from that of the ambient droplets. We named these droplets “spongelets” because they had medium levels for most ADTs without any enrichment for specific ADTs compared to true cell clusters. Although our analysis did not reveal the source of the spongelets, we hypothesize that they could be due to the presence of debris from the dissociation procedure or dying cells with a permeable membrane. Further work will be required to identify the source of this artifact and understand how experimental parameters can be modified to decrease the contribution of spongelets to the background.

These observations motivated us to design a decontamination algorithm that decomposes the ADT count matrix into three components: 1) native counts representing the contribution from true cells, 2) ambient counts representing the contribution from ambient material, and 3) background counts representing the contribution from other low level contamination sources such as nonspecific binding or spongelets. Importantly, this model allows DecontPro to work in situations where the raw empty droplet matrix is not available which is often the case in public repositories. To estimate the ambient profile of a dataset when the empty drop matrix is not available, we use the average of the true cells. We found that the average of the true cells was highly correlated to the ambient droplets in three out of the four datasets examined. The lower correlation in the Golomb dataset was primarily due to one ADT (Ly6C) which had lower than expected levels in the cells given the level in the ambient droplets. Other experimental procedures such as cell sorting for particular populations may also break the assumption that the cell average will accurately approximate the profile of the ambient droplets. In these cases, our software allows users to input the empty droplets to more accurately estimate the ambient profile.

Overall, DecontPro can be used as an important quality assessment tool that estimates the levels of different sources contributing to the contamination in ADT data. The computational decontamination of ADT counts with DecontPro will aid in downstream clustering and visualization and can be systematically included in analysis workflows.

## Methods

### Empty droplet analyses

The PBMC 10k dataset containing a total of 6,794,880 raw droplets and 7,865 filtered droplets was downloaded from the 10x Genomics website^21^. Droplets with zero total counts for both ADT and RNA were filtered. The library sizes for ADT and RNA were calculated by summing all the ADT and RNA counts in each droplet. The droplets with barcodes in the raw but not filtered dataset were named empty droplets. Clusters B, C and D in empty droplets were identified using k-means with the number of clusters set to 3. Profiles of cluster C (spongelet) and D (ambient) were calculated by summing ADT counts over droplets and normalizing the vector to one. The cell background profile was calculated by summing ADT counts over droplets that fall into the background peak and normalizing. The background peaks are manually identified using the bounds shown in **Supplementary Figures S14-S16**. Cell clusters were identified with markers: MonoCD14: (CD14+), CD4TMemory: (CD3+, CD4+, CD45RO+), CD4TNative: (CD3+, CD4+, CD45RA+), CD8TMemory: (CD3+, CD8+, CD45RO+), NK: (CD16+, CD56+), B: (CD19+), CD8TNaive: (CD3+, CD8+, CD45RA+).

### Data Generation Model

For a droplet *i*. containing a cell of type *k_i_* having the library size *L_i_*, we assume the *j^th^* ADT count *x_ij_* is a realization from a Poisson distribution with rate parameter *λ_ij_*:

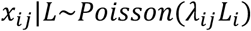

When there is a total of *I* droplets, *J* ADTs and *K* cell types, the count matrix can be organized into an *I* by *J* matrix *X* = [*x_ij_*] and a corresponding cell type indicator vector *k* = [*k_i_*]. We further assume that the overall expected rate of counts for an ADT in a cell (*λ_ij_*) can be described as a sum of three components:

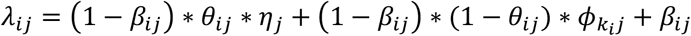

where *ϕ_k_i_j_* is the normalized rate of counts for ADT *j* in cell population *k_i_*, *η_j_* is the normalized rate of counts due to the ambient source for ADT *j, θ_ij_* is the proportion of ambient material for ADT *j* in cell *i*, and *β_ij_* is the normalized rate of contamination from all other background sources including spongelets and non-specific binding.

We assume that some level of general background (*β_ij_*) will be present for each ADT in each cell and that the rates of native or ambient ADTs can be quantified in each droplet after subtracting out the background rate (1 – *β_ij_*). More specifically, the normalized rate of ambient ADTs can be expressed as the expected proportion of counts coming from the ambient distribution times the level of ambient contamination in that cell after excluding the general background rate: (1 – *β_ij_*) * *θ_ij_* * *η_j_*. The rate of true native ADT counts for a given cell can be expressed as the expected number of counts from that cell type after excluding the proportion of counts from both ambient contamination and background noise: (1 – *β_ij_*) * (1 – *θ_ij_*) * *ϕ_k_i_j_*.

We put prior distributions on the rates of ambient contamination for all ADTs in a droplet and the background contamination for each ADT across all cells. *β_ij_* is drawn from a truncated normal parameterized by mean *μ_j_* and bounded by (0, 0.5) while *θ_ij_* is drawn from a truncated normal parameterized by mean *δ_i_* and bounded by (0,1):

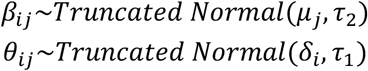

*τ_1_* and *τ_2_* are hyperparameters that control how strongly individual contamination rates can deviate from the prior mean rates of *δ_i_* and *μ_j_*, respectively. The plate diagram for the model is shown below.

### Application of DecontPro to ADT datasets

Datasets were preprocessed by filtering out cell droplets with top and bottom one percent of ADT and RNA total library sizes, and droplets with 15% or higher mitochondrial gene counts. Cell clusters were generated using the Seurat package. The prior parameters were set to *τ*_1_(PBMC 10k: 2e-5, PBMC 5k: 5e-5, MALT 10k: 4e-5, Golomb: 2e-5) and *τ*_2_(PBMC 10k: 2e-6, PBMC 5k: 5e-6, MALT 10k: 2e-6, Golomb: 2e-6). The inference was done using the variational inference implementation in Stan^22^. *θ_ij_* was initialized 1e-4 and *β_ij_* was initialized 1e-2. The max number of iterations were set to 50,000.

### Benchmarking of decontamination tools

Four datasets were used in benchmarking. PBMC 10k: PBMCs from a healthy donor stained with 17 Total-Seq-B antibodies (7,865 filtered droplets). PBMC 5k: PBMCs from a healthy donor stained with 31 Total-Seq-B antibodies (5,527 filtered droplets). MALT 10k: Dissociated Extranodal Marginal Zone B-Cell Tumor cells stained with 17 Total-Seq-B antibodies (8,412 filtered droplets). These three datasets were downloaded from 10x Genomics website. Golomb: Brain samples of mice with antibiotics induced gut microbiota depletion (30,569 filtered droplets). A panel of 31 antibodies was used for staining the samples of 3 young mice and 3 aged mice, and the samples were pooled together using hashtag oligo.

The clustering of datasets was generated using the Seurat package. Silhouette widths were calculated on datasets after CLR normalization and averaged by clusters. The positive and negative scores for each dataset were calculated by finding the percentage of droplets in the dataset that belong to the clusters and have the markers expressed more than the threshold after decontamination. The threshold was set to 1 except when evaluating dsb, we also used a high threshold of 5 (reported the results as “dsb (high threshold)” in the main manuscript). The cell clusters and markers used to calculate the scores are listed in **Supplementary Table S1**.

## Acknowledgements

We thank Dr. Siyuan Zhang for providing access to the Golomb dataset. We also thank Samantha Golomb, Yusuke Koga, Junxiang Xu, and Charley Xi for their feedback and help. This work was funded by the National Library of Medicine (NLM) R01LM013154-01 (JDC, MY) and the Chan Zuckerberg Initiative DAF Data Insights grant “Methods and software for decontamination of single-cell data”.

## Availability of data and materials

The PBMC 10k^23^, PBMC 5k^24^ and MALT 10k^25^ datasets used to support the study were downloaded from 10x Genomics. Golomb dataset was from GEO: GSE148127^26^ and the corresponding author of the study.

DecontPro is freely available as an R package on GitHub: https://github.com/campbio/decontX.

## Notes

https://github.com/campbio/decontX

### Competing Interest Statement

The authors have declared no competing interest.

### Summary of Updates

Proofread the manuscript and updated figures. Added GitHub link to the software.

