## Supplementary for "Characterization and decontamination of background noise in droplet-based single-cell protein expression data with DecontPro"

**Table S1. Markers used to calculate positive and negative scores.**

| Dataset | Cell Cluster | Scores | Marker |
|---|---|---|---|
| **PBMC 10K** | CD14 Monocyte | + | CD14 |
| | CD4 naïve T-cell | + | CD4, CD45RA |
| | | - | CD8a, CD45RO |
| | CD4 memory T-cell | + | CD4, CD45RO |
| | | - | CD8a, CD45RA |
| | NK | + | CD16, CD56 |
| | CD8 memory T-cell | + | CD8a, CD45RO |
| | | - | CD4, CD45RA |
| | B | + | CD19 |
| | | - | CD3 |
| | CD8 naïve T-cell | + | CD8a, CD45RA |
| | | - | CD4, CD45RO |
| | CD16 monocyte | + | CD16 |
| **PBMC 5K** | CD14 Monocyte | + | CD14 |
| | CD4 naïve T-cell | + | CD4, CD45RA |
| | | - | CD8a, CD45RO |
| | CD4 memory T-cell | + | CD4, CD45RO |
| | | - | CD8a, CD45RA |
| | NK | + | CD16, CD56 |
| | CD8 memory T-cell | + | CD8a, CD45RO |
| | | - | CD4, CD45RA |
| | B | + | CD19 |
| | | - | CD3 |
| **MALT 10K** | CD4 memory T-cell | + | CD4, CD45RO |
| | | - | CD8a, CD45RA |
| | CD8 memory T-cell | + | CD8a, CD45RO |
| | | - | CD4, CD45RA |
| | B | + | CD19 |
| | | - | CD3 |
| **Golomb** | Microglia | + | CD11b |
| | | - | CD3, CD8a |
| | CD8 T-cell | + | CD3, CD8a |
| | | - | CD45R-B220 |
| | B | + | CD45R-B220 |
| | | - | CD3, CD8a |

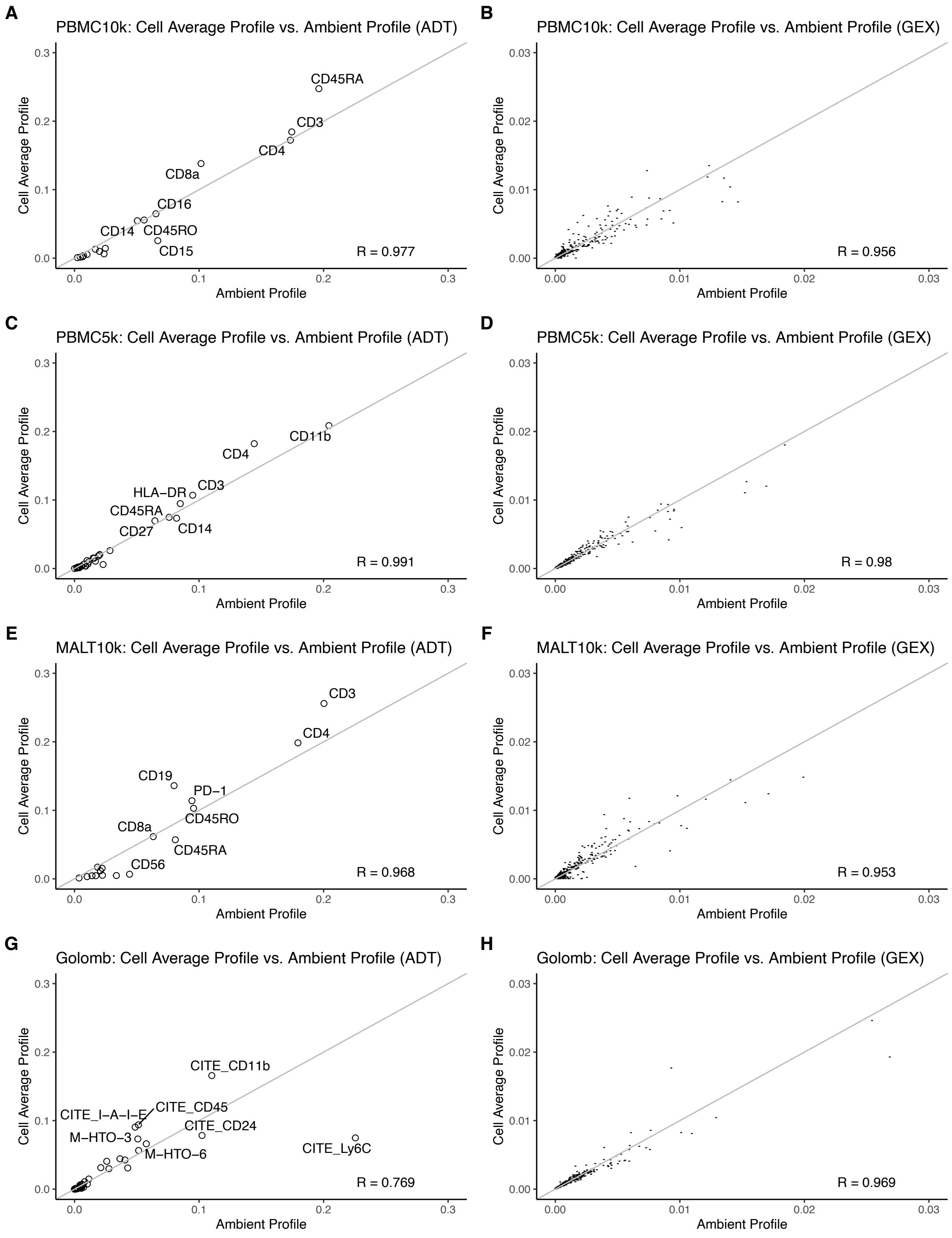


**Figure S1. Correlation between Cell profile and Ambient profile.** The average cell profile of ADTs (**A. C. E. G.**) from the Cell droplets was found to be highly correlated with the ambient profile which represents the ambient signal. This finding was reproduced using the gene expression data of the droplets (**B. D. F. H.**).

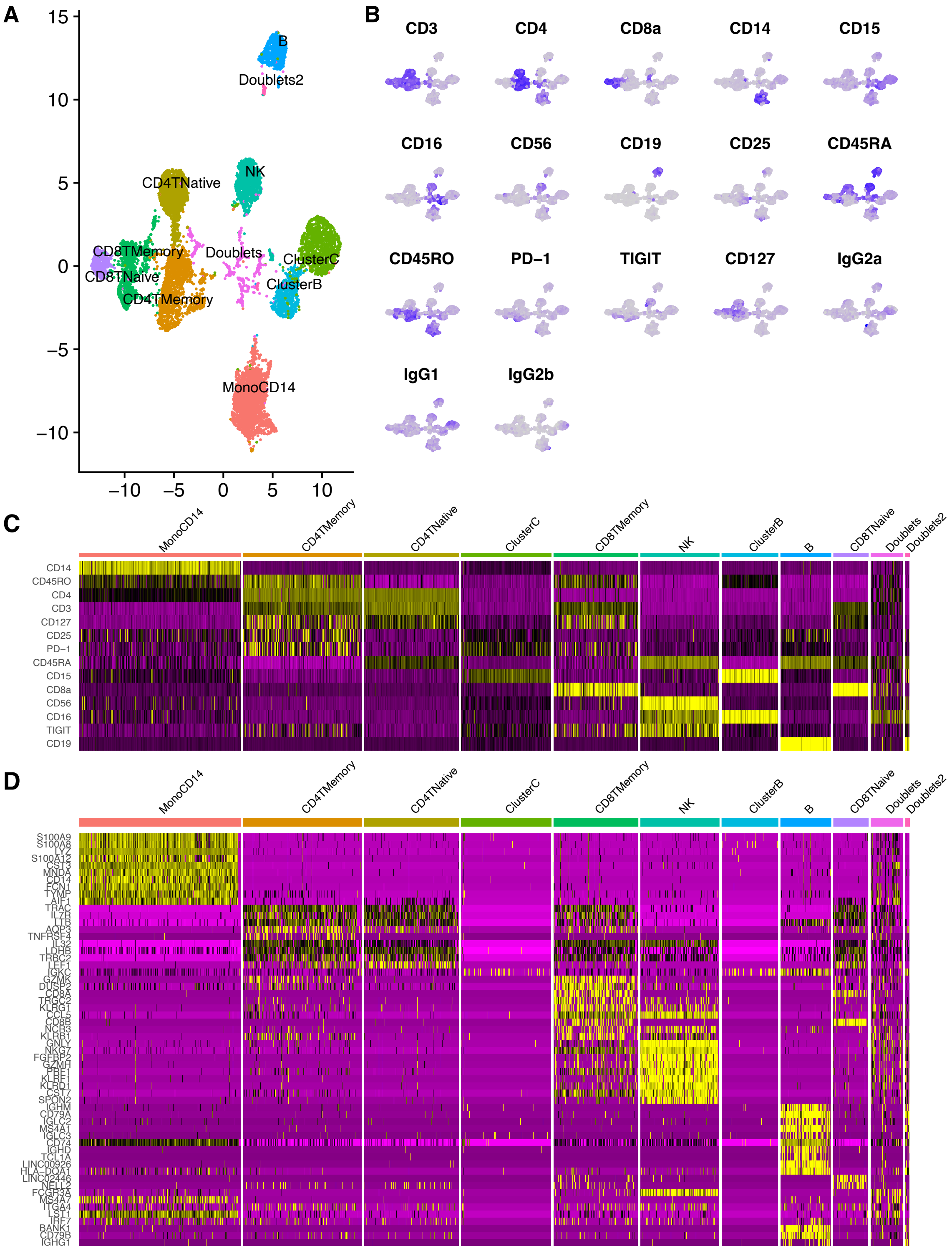


**Figure S2. Annotating PBMC 10K dataset. A.** UMAP of droplets from the Cell, Cluster B and Cluster C. One thousand droplets were each randomly sampled from the Cluster B and C to join the Cell cluster for clustering. **B.** Feature plot of ADTs on the UMAP. **C.** Differential expression analysis for each cluster using the ADT expression data. **D.** Differential expression analysis for each cluster using the gene expression data.

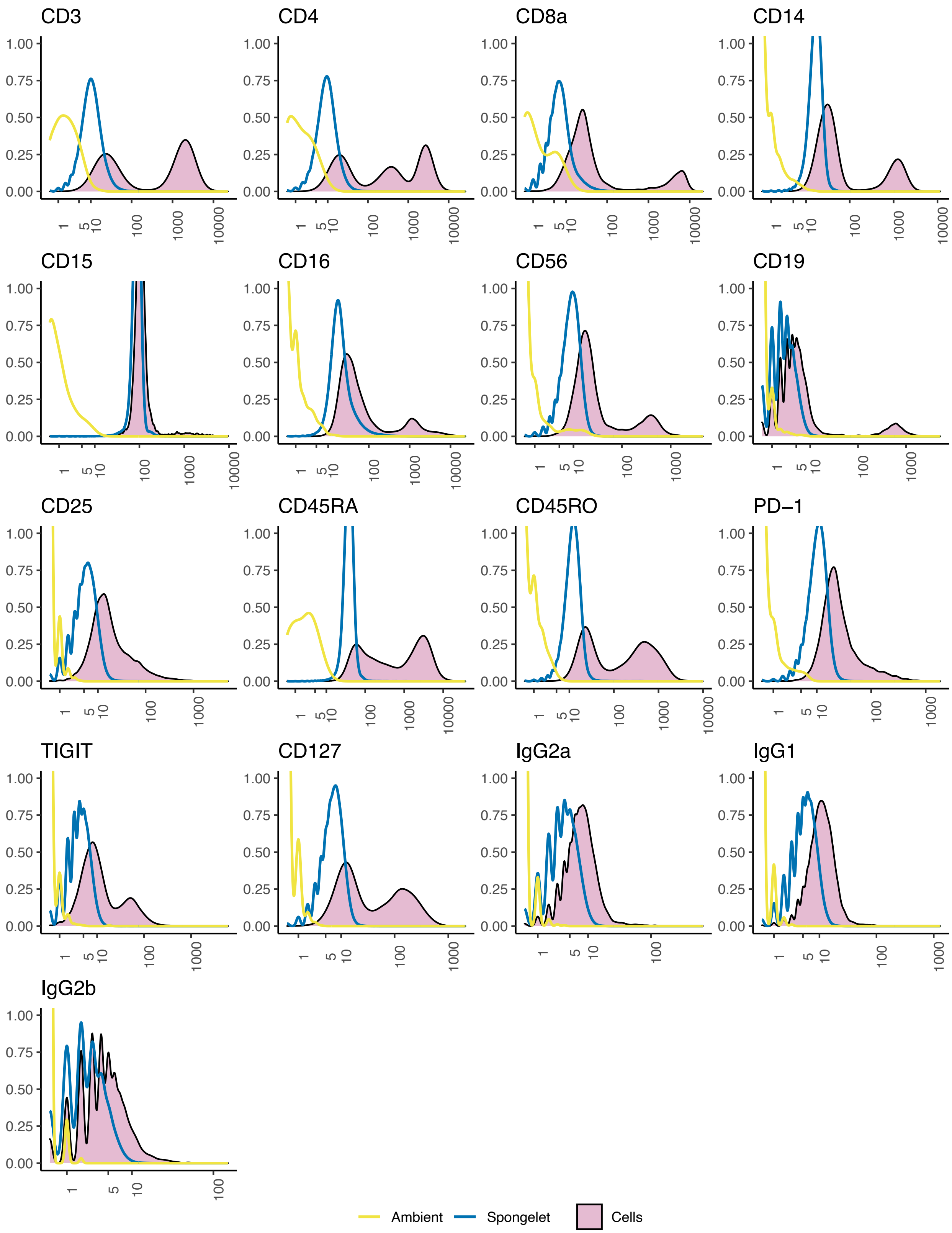


**Figure S3. Density plot for ADTs in PBMC 10K dataset.** Density plots of all ADTs in the dataset for droplets from the Cell, Spongelet and Ambient clusters.

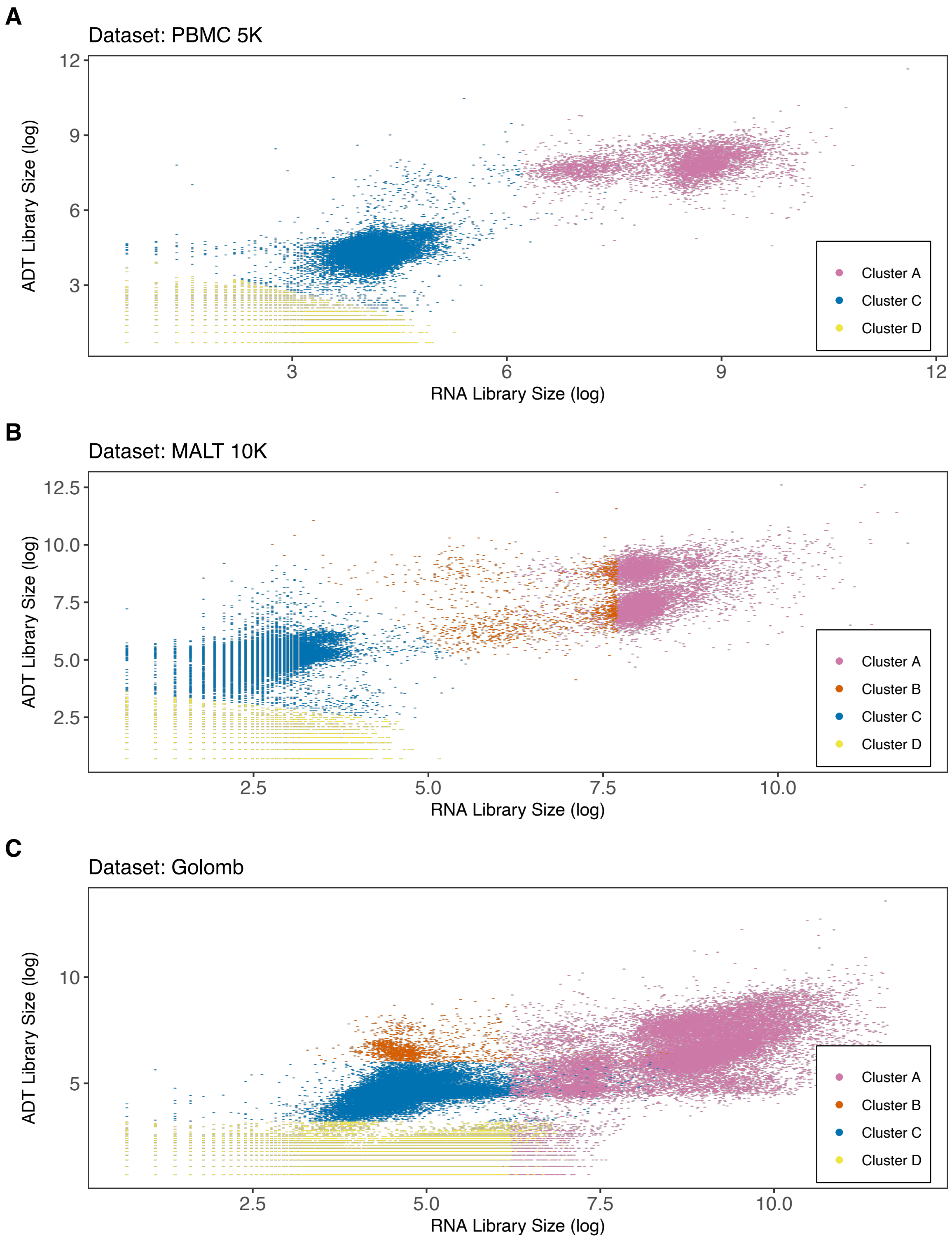


**Figure S4. Identifying clusters in empty droplets for three datasets.** Droplets after Cell Ranger filtering are labeled Cell, and the rest are the empty droplets. We identified clusters in the empty droplets using k-means for all datasets, except Golomb dataset where clusters were manually identified based on empty droplets' ADT library sizes. The Cluster C and Cluster D were used for downstream contamination analysis.

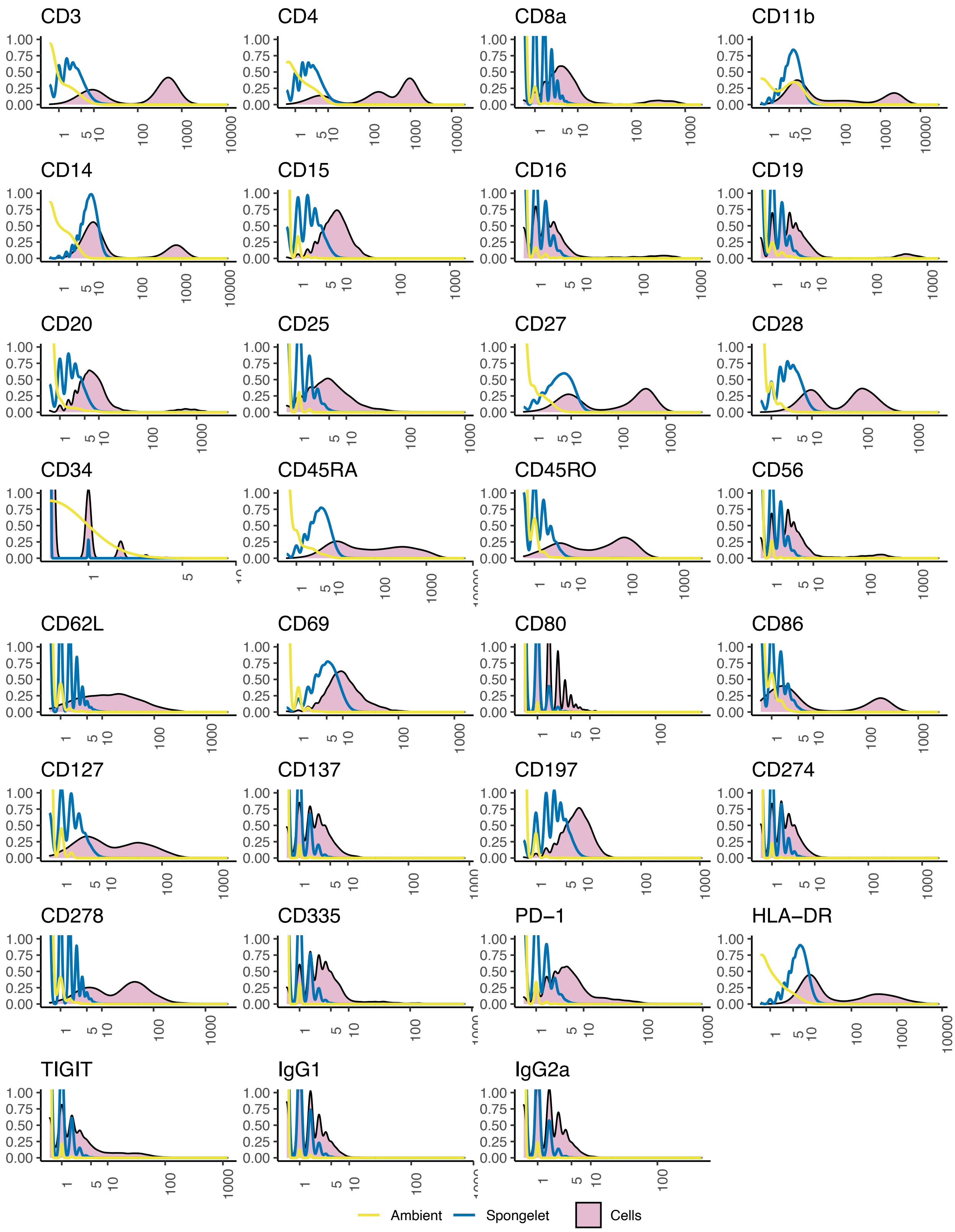


**Figure S5. Density plot for ADTs in PBMC 5K dataset.** Density plots of all ADTs in the dataset for droplets from the Cell, Spongelet and Ambient clusters.

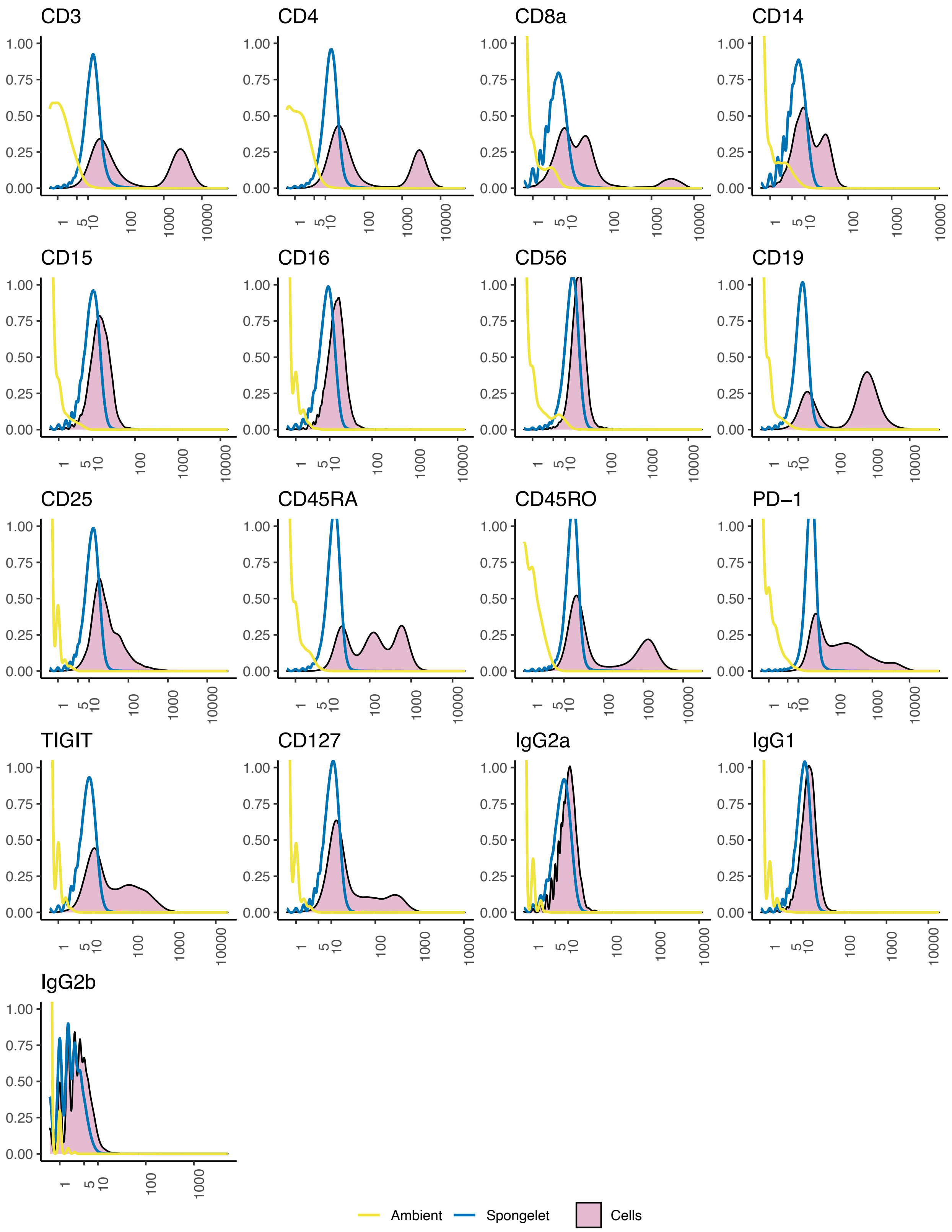


**Figure S6. Density plot for ADTs in MALT 10K dataset.** Density plots of all ADTs in the dataset for droplets from the Cell, Spongelet and Ambient clusters.

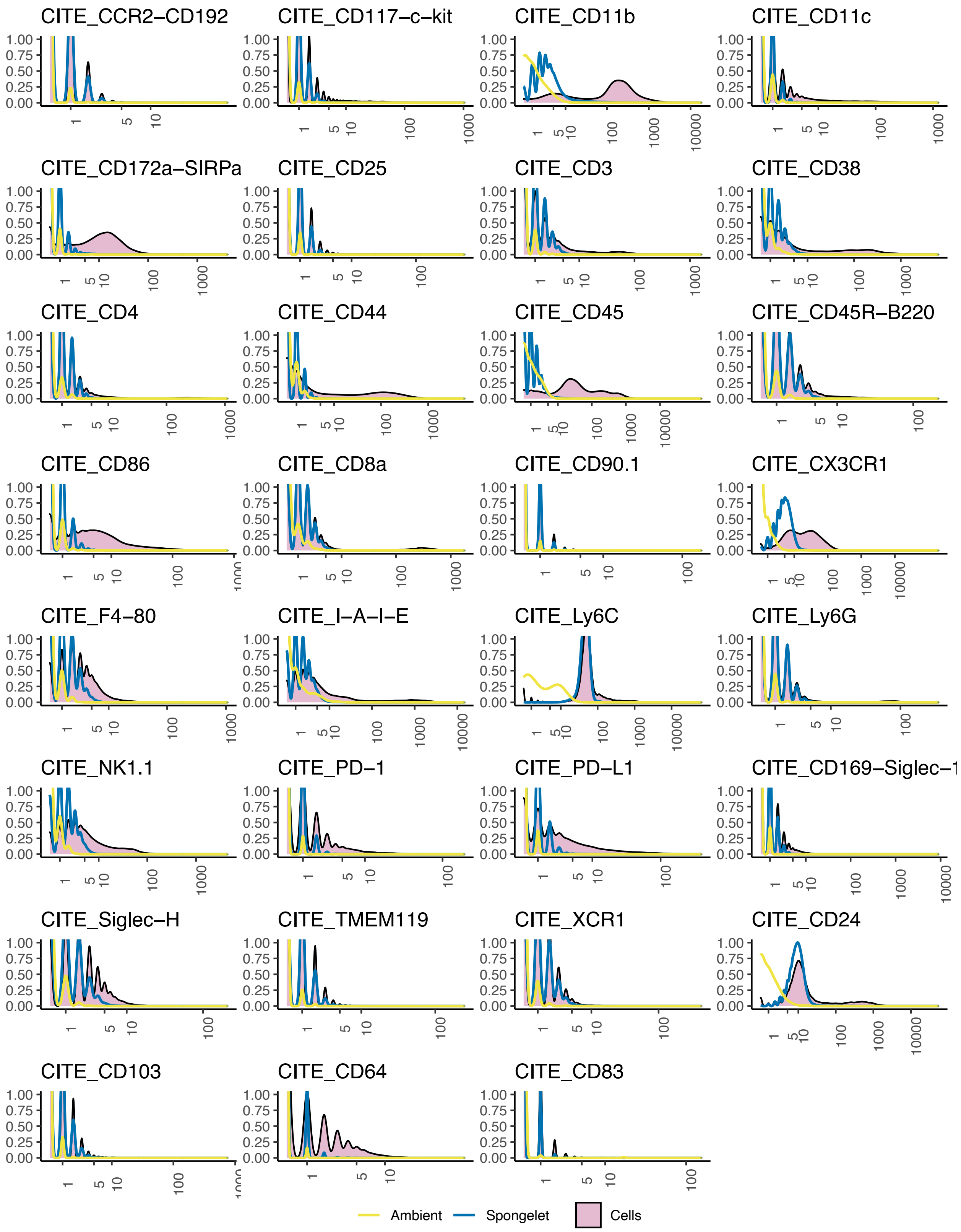


**Figure S7. Density plot for ADTs in Golomb dataset.** Density plots of all ADTs in the dataset for droplets from the Cell, Spongelet and Ambient clusters.

**A**

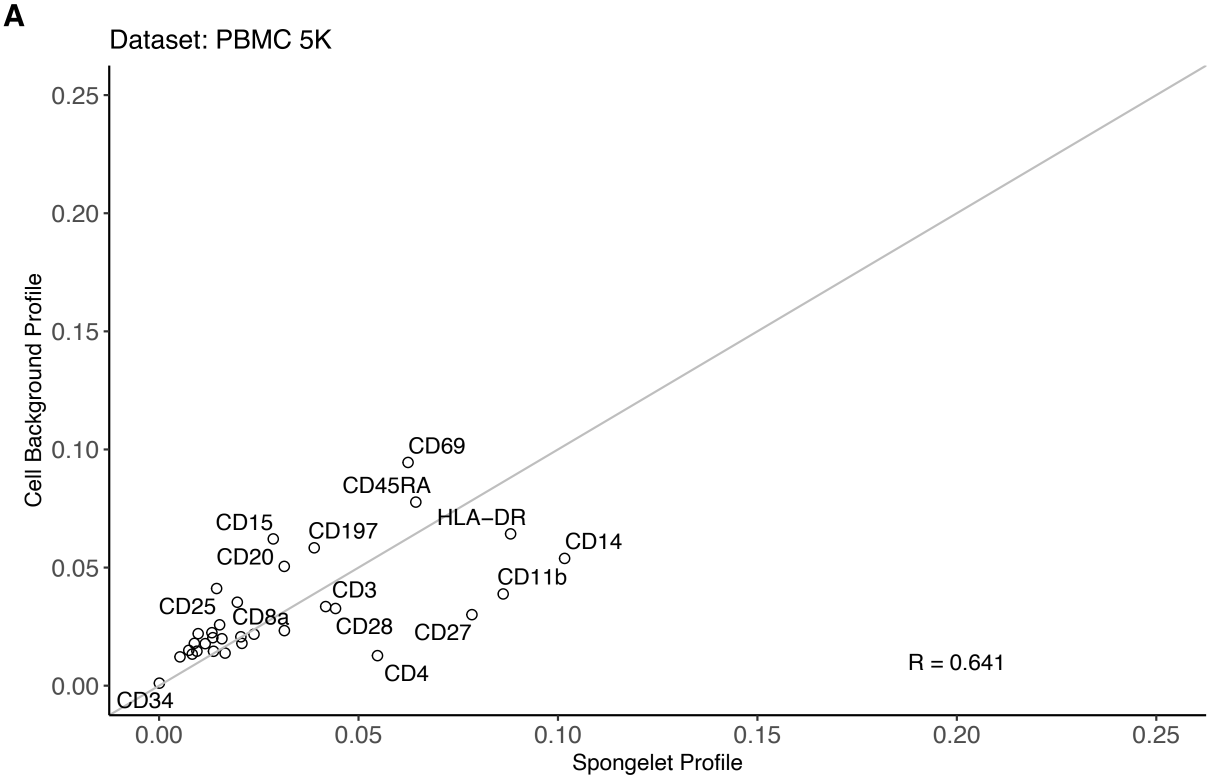


**B**

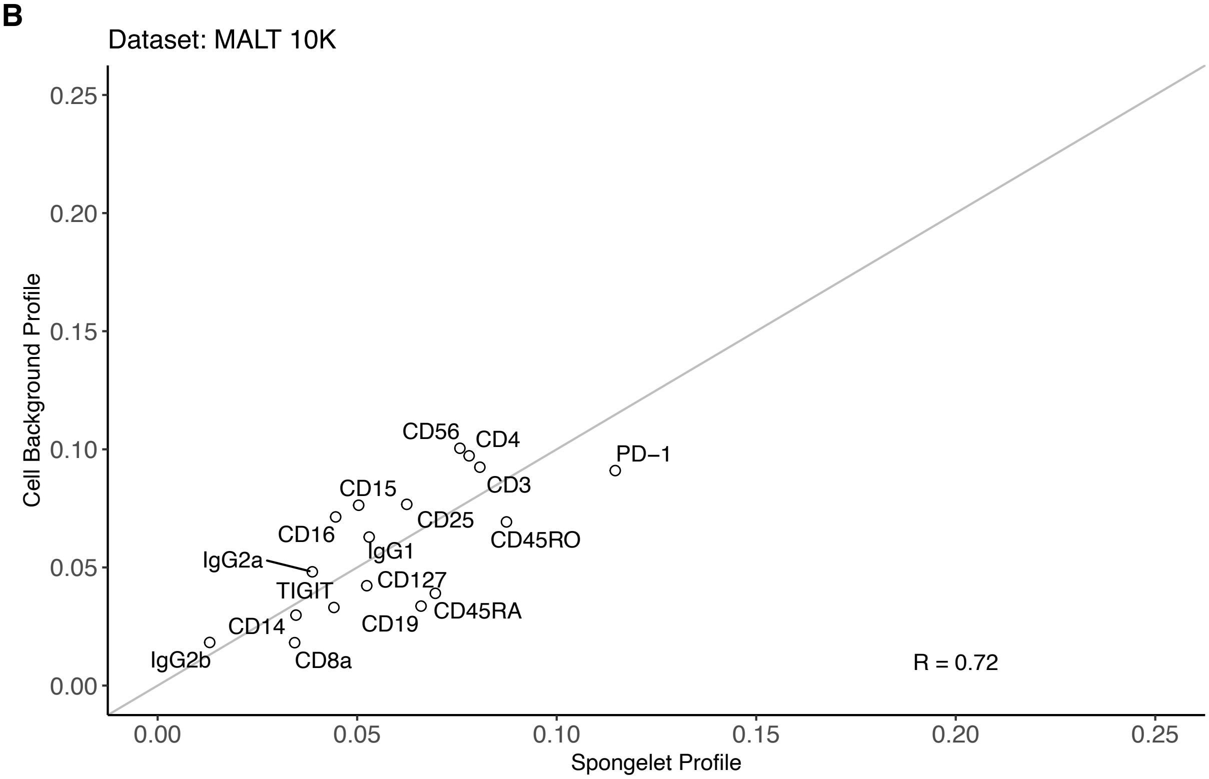


**Figure S8. Correlation between Cell background profile and Spongelet profile** for **A.** PBMC 5K dataset and **B.** MALT 10K dataset.

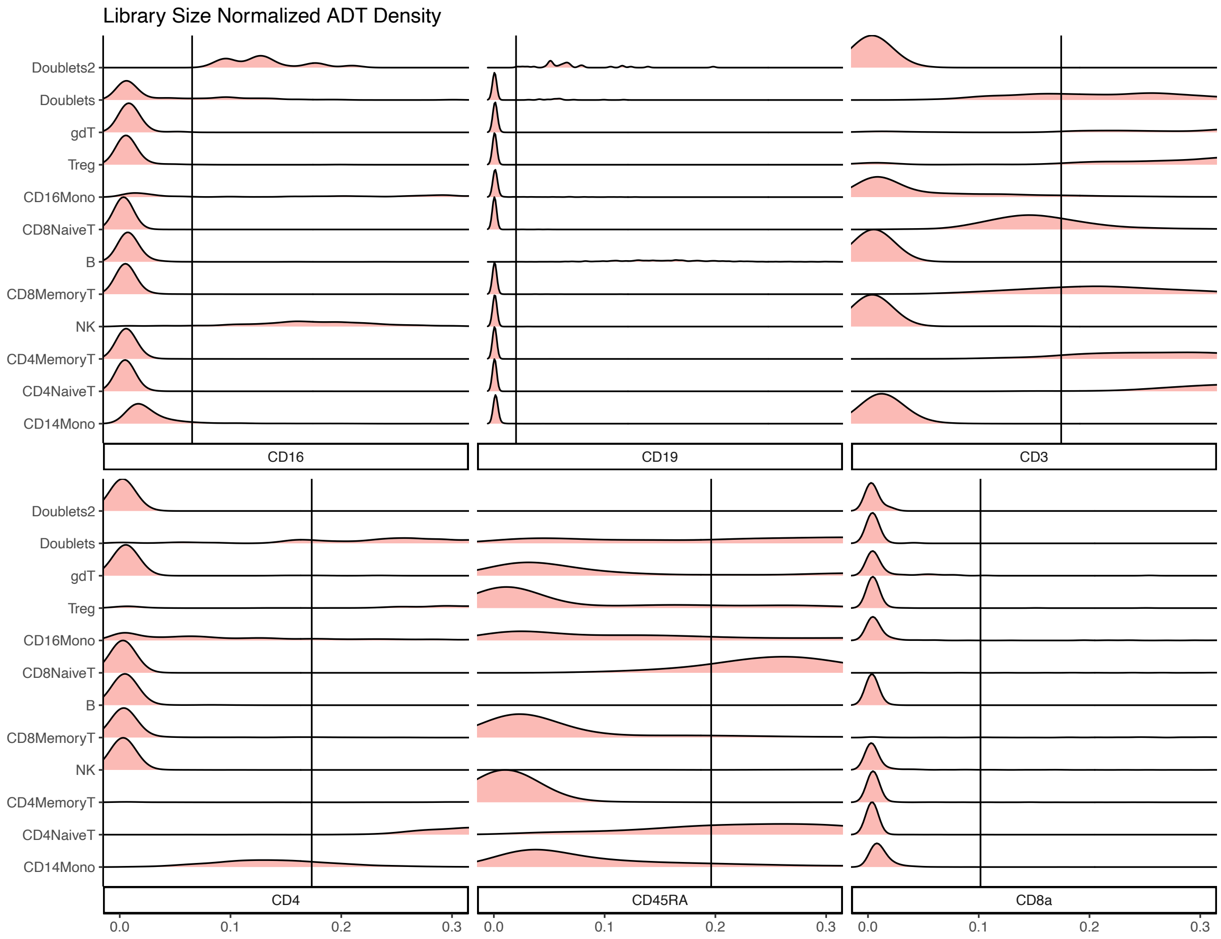


**Figure S9. Density plots for library size-normalized droplets for PBMC 10K dataset.** ADTs not expected to be expressed in cell types show a peak at a near zero level, in contrast to a more spread-out peak at a higher level when they are natively expressed in other cell types. When overlapping the normalized ambient profile on top of the density plot, the ambient distribution differs from the mean of the margin peaks.

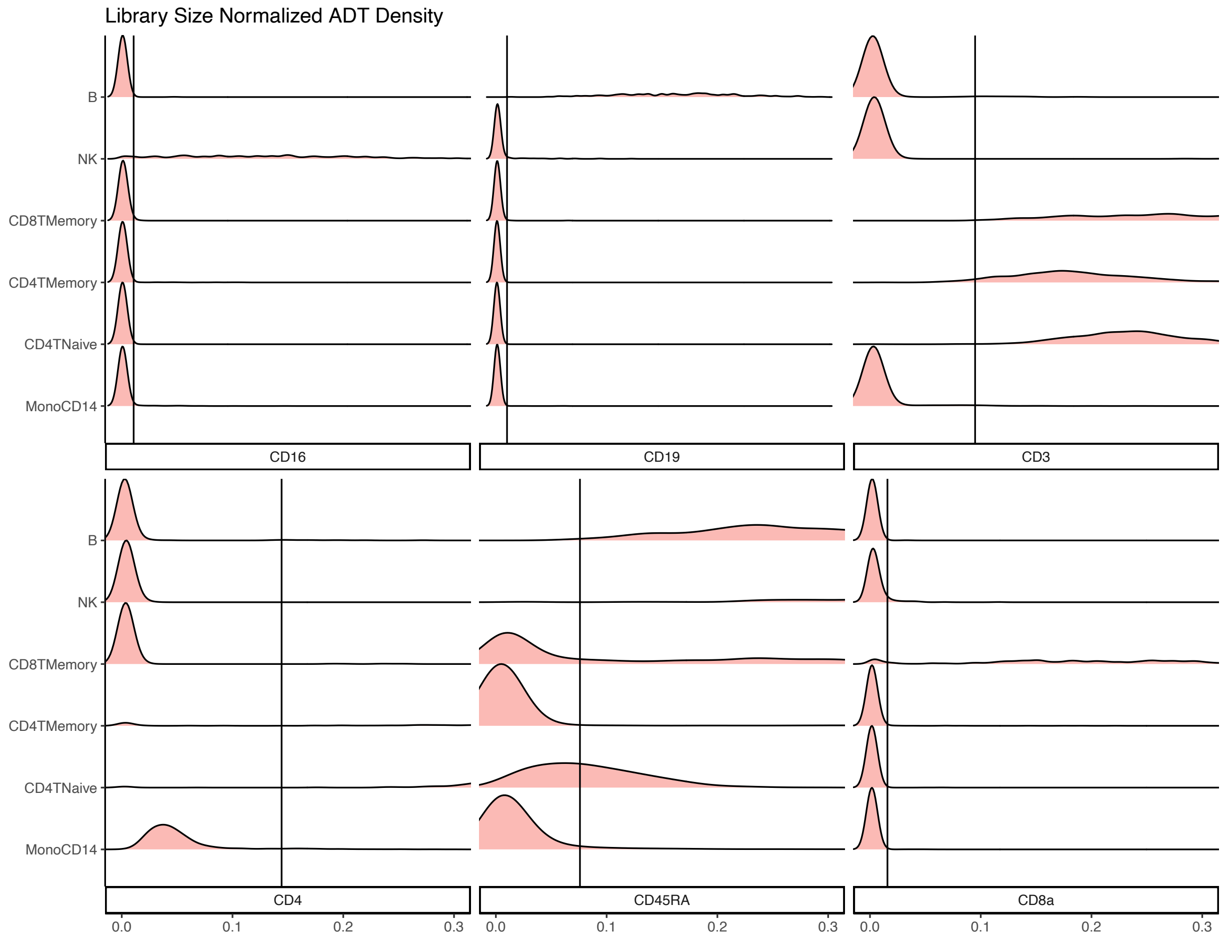


**Figure S10. Density plots for library size-normalized droplets for PBMC 5K dataset.** ADTs not expected to be expressed in cell types show a peak at a near zero level, in contrast to a more spread-out peak at a higher level when they are natively expressed in other cell types. When overlapping the normalized ambient profile on top of the density plot, the ambient distribution differs from the mean of the margin peaks.

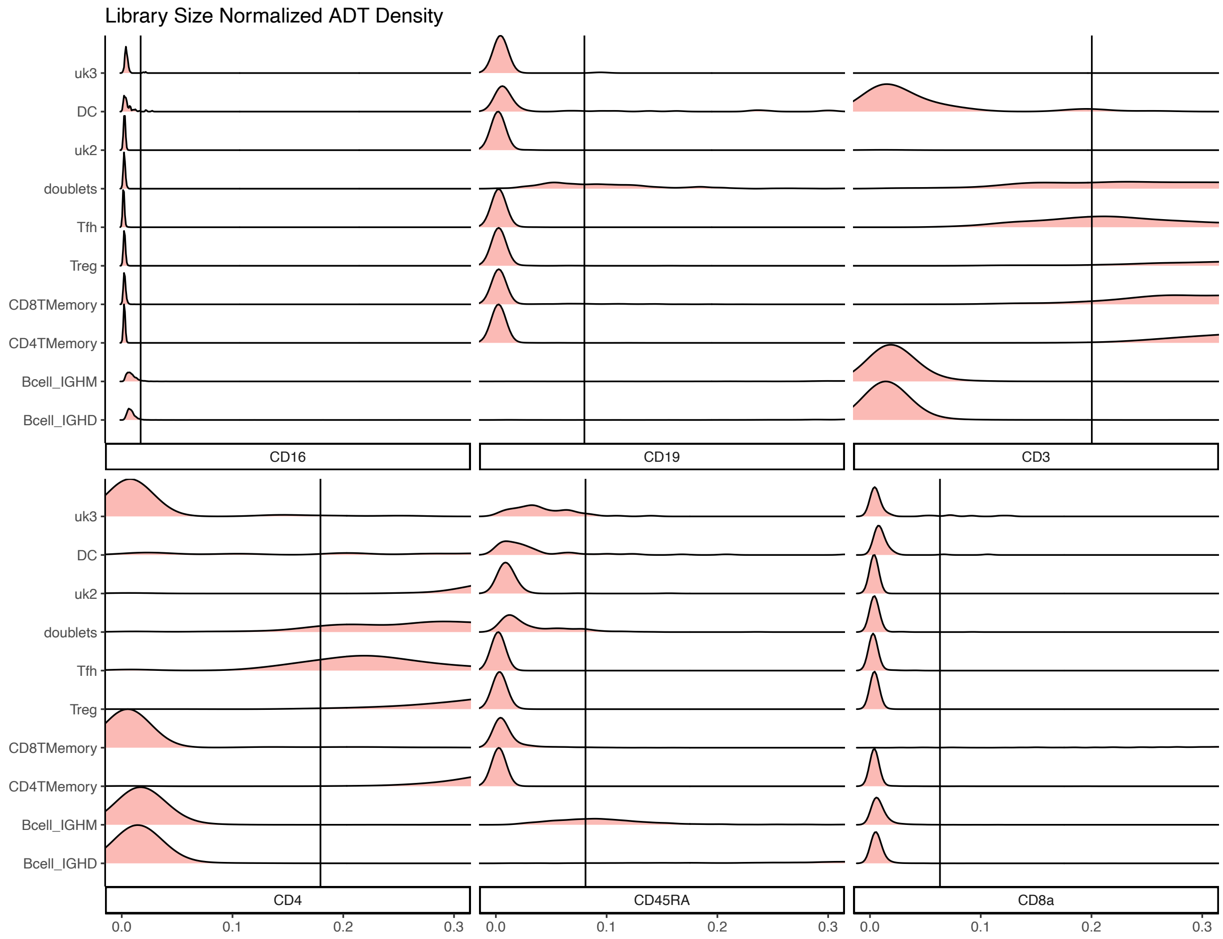


**Figure S11. Density plots for library size-normalized droplets for MALT 10K dataset.** ADTs not expected to be expressed in cell types show a peak at a near zero level, in contrast to a more spread-out peak at a higher level when they are natively expressed in other cell types. When overlapping the normalized ambient profile on top of the density plot, the ambient distribution differs from the mean of the margin peaks.

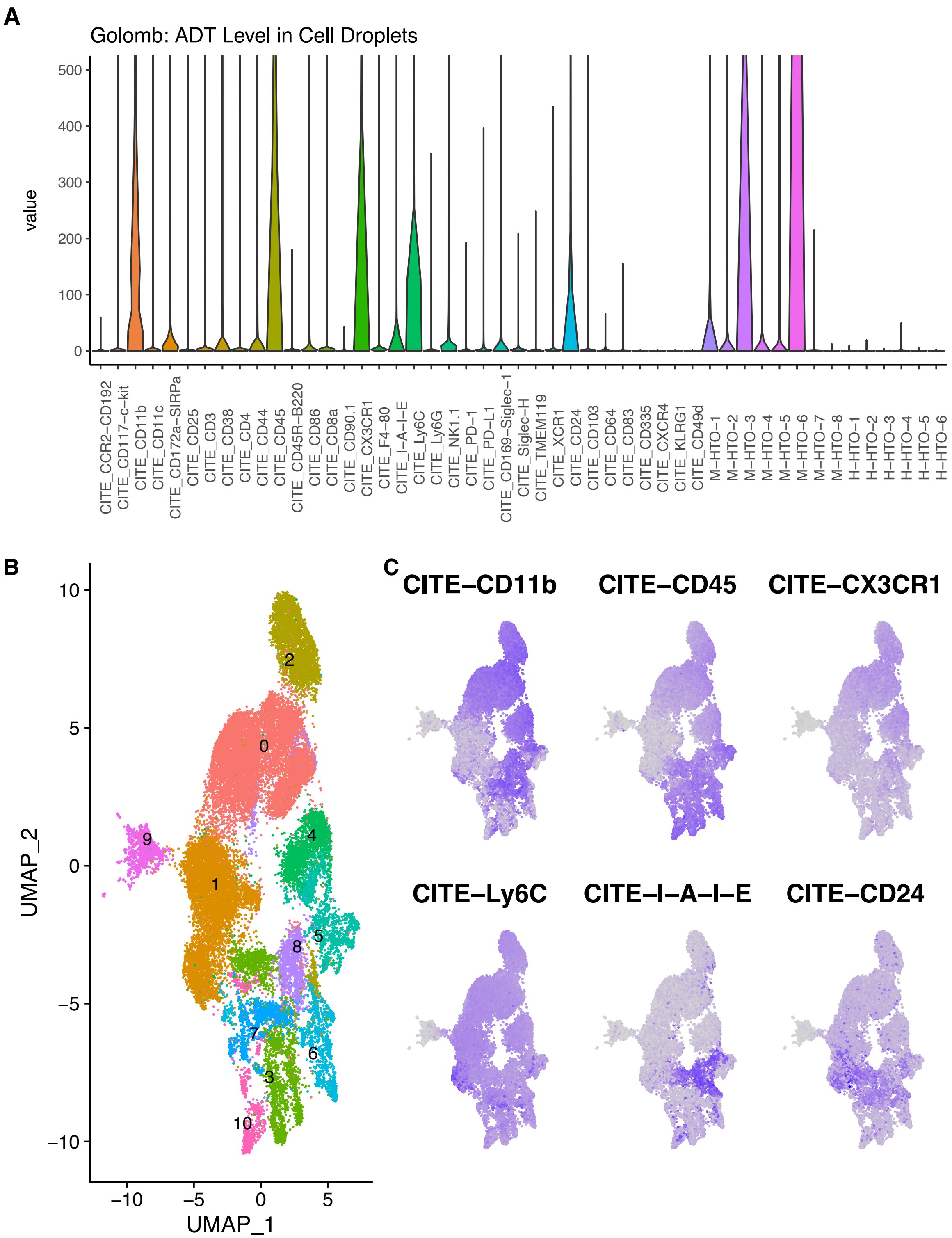


**Figure S12. Ly6C as an outlier ADT among the highly stained ADTs in Golomb dataset. A.** Violin plot of ADT expression level among the cell droplets. Ly6C is one of the highly stained ADTs in the panel. **B.C.** Highly stained ADTs identified in **A** were shown on a UMAP. Ly6C shows high expression globally, in contrast to other ADTs with localized high expression.

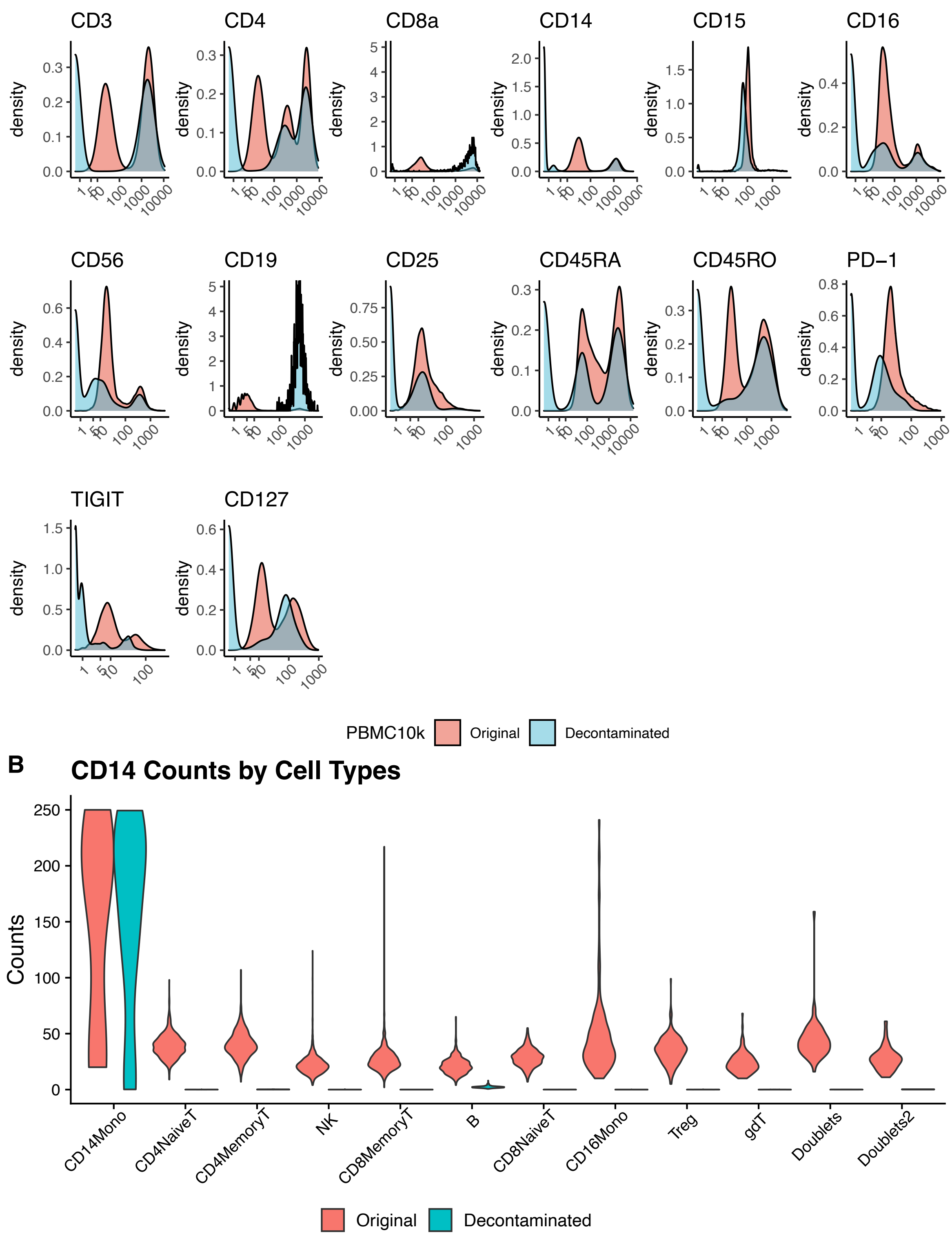


**Figure S13. Comparison of uncorrected original counts and decontaminated counts of PBMC 10K dataset after DecontPro. A.** Density of each ADT expression before and after decontamination shows the background peak is largely reduced. **B.** CD14 counts are greatly reduced in clusters other than CD14 Monocyte cluster.

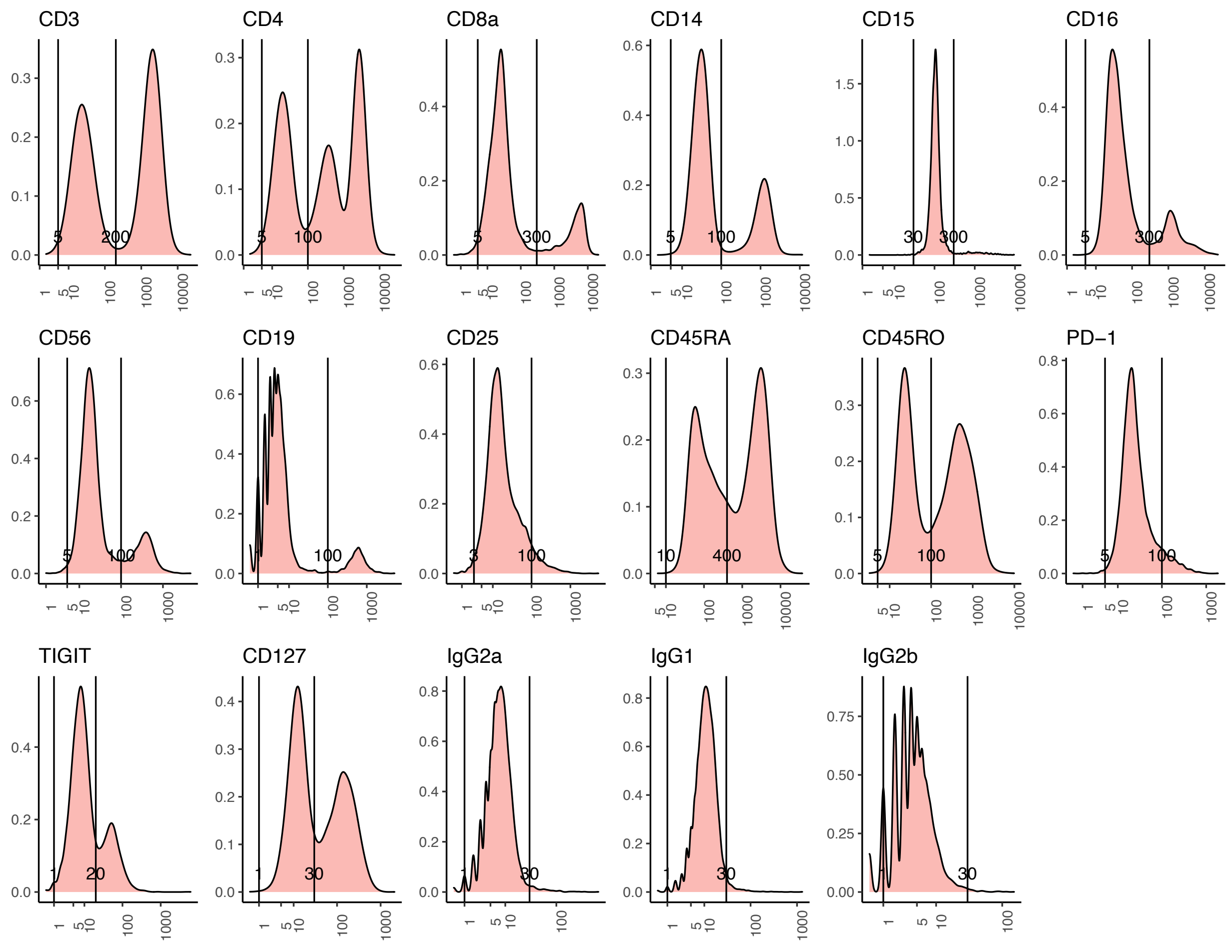


**Figure S14. Identifying background peaks for PBMC 10K dataset.** Background peaks for ADTs were identified and labeled using an upper bound and a lower bound.

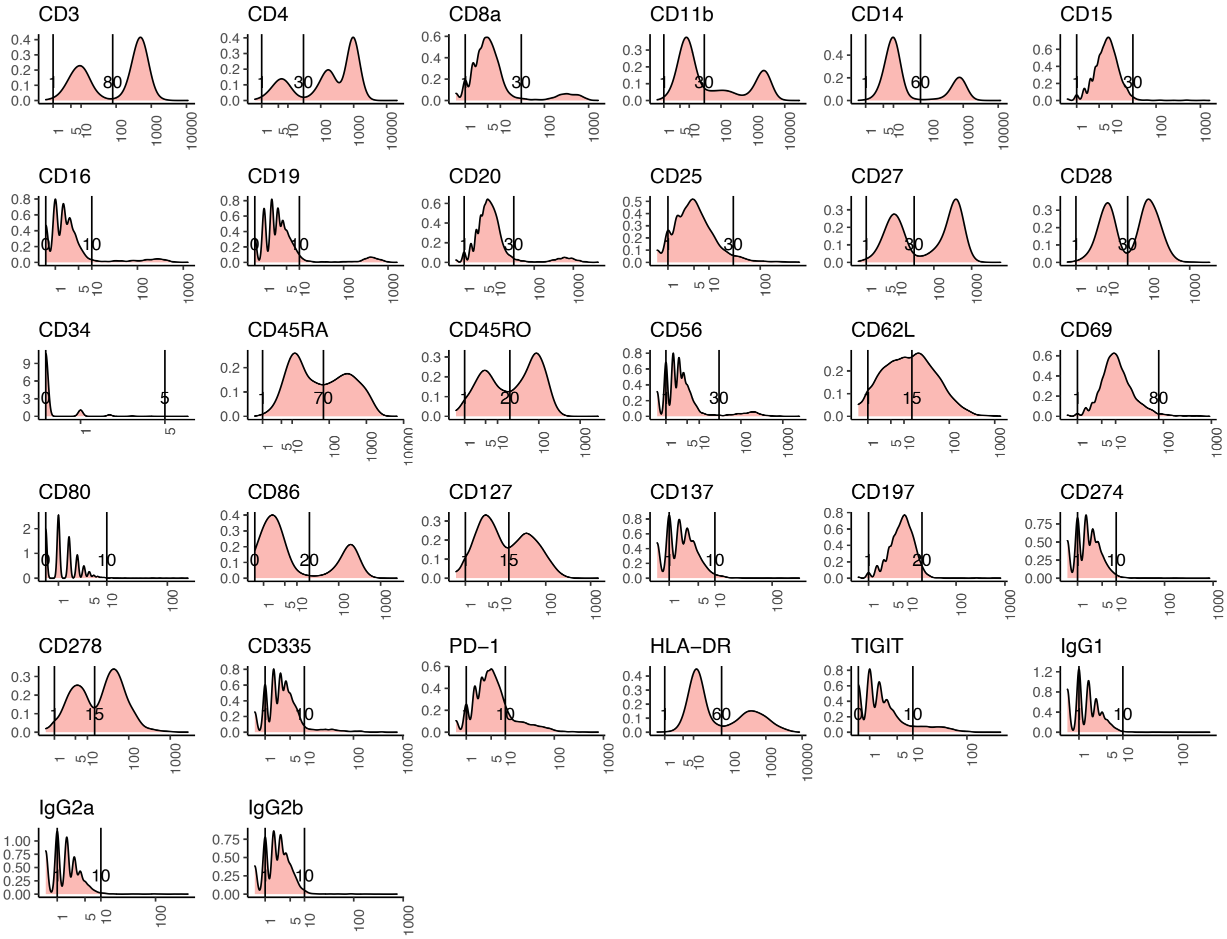


**Figure S15. Identifying background peaks for PBMC 5K dataset.** Background peaks for ADTs were identified and labeled using an upper bound and a lower bound.

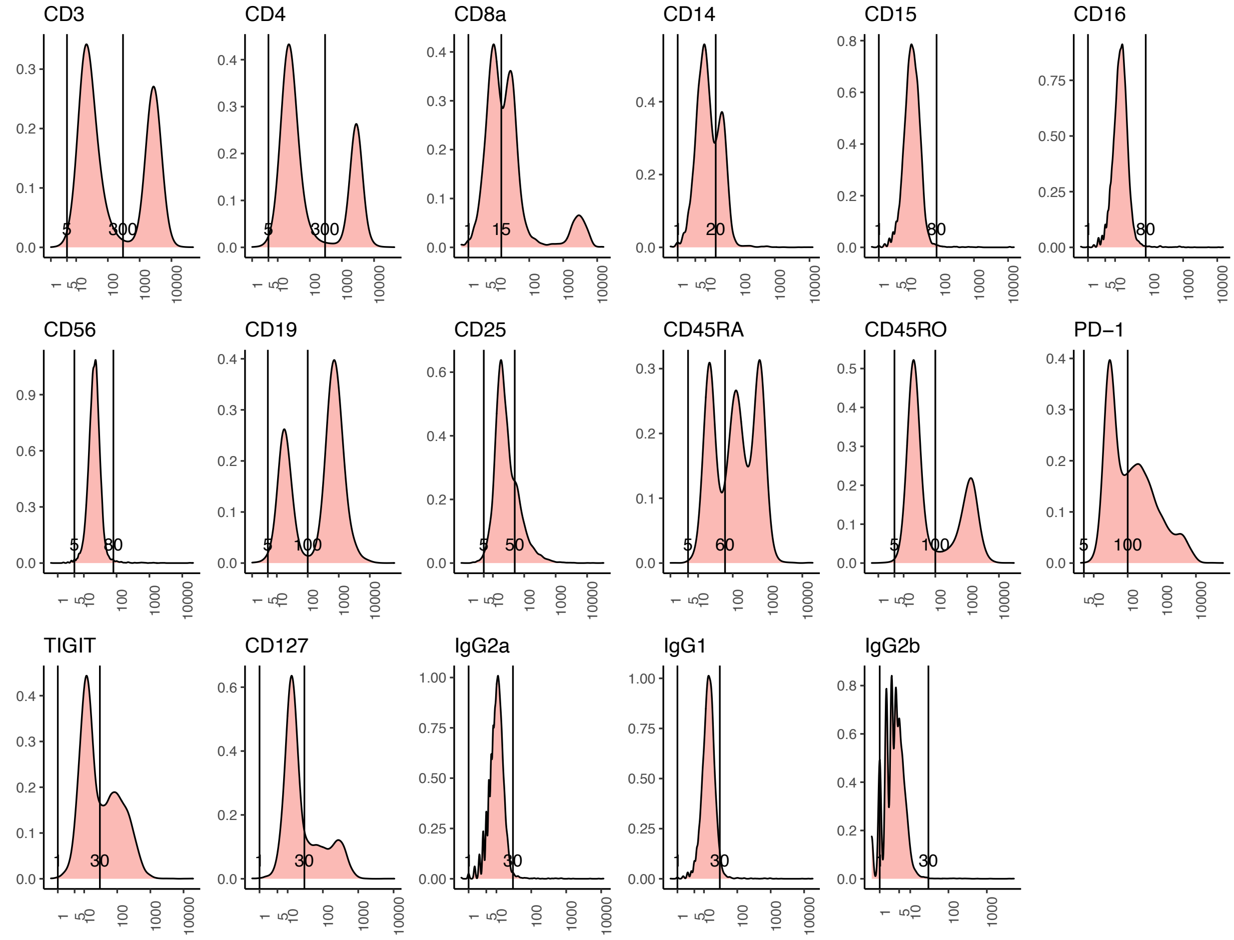


**Figure S16. Identifying background peaks for MALT 10K dataset.** Background peaks for ADTs were identified and labeled using an upper bound and a lower bound.
